# Programmable multiplexed phosphosignaling networks in bacteria

**DOI:** 10.1101/2025.09.09.674673

**Authors:** Xinyue Song, Bentley Lim, Mimi C. Yung, Shangri-La Hou, Yongqin Jiao, W. Seth Childers

## Abstract

Natural bacterial two-component systems use crosstalk between histidine kinases to integrate multiple environmental cues, enabling complex multi-signal decisions. In comparison, most synthetic bacterial phosphosignaling circuits remain confined to single-input designs. Here, we establish a bottom-up framework for phosphosignaling circuits using two histidine kinases (2HKs) that converge on a shared response regulator (RR). As a proof of concept, we built a synthetic 2HK–1RR circuit in *Escherichia coli* that consists of a blue-light sensor (YF1) and an indole-3-aldehyde sensor, integrating opto- and chemo-signals through their shared RR. By using distinct dimerization tags on each HK, tuning HK activities, and reversing HK response modes, we program a spectrum of logic behaviors. These include ternary gates (an inverse count gate, a count gate, and two context gates), as well as a binary NAND gate. This work establishes fundamental phosphosignaling circuit design rules based on two-component systems that expand the scope of synthetic gene circuit design and pave the way towards complex, multi-input environmental biosensors and programmable therapeutics.

## INTRODUCTION

Bioengineering increasingly requires synthetic circuits capable of integrating multiple environmental inputs to reduce off-target effects in therapeutic applications,^1^ control microbial consortia,^2^ and enhance specificity in environmental monitoring.^3, 4^ In natural bacteria, a key mechanism enabling such complex decision-making is through crosstalk, such as interactions where components from distinct signaling pathways converge on shared targets. In bacterial two-component systems (TCSs),^5^ crosstalk among many histidine kinases (HKs)^6^ is common in developmental processes such as sporulation,^7^ asymmetric cell division,^8, 9^ and biofilm formation.^10^ For example, in *Caulobacter crescentus*, DivJ functions as a kinase and PleC as a phosphatase to oppositely regulate the phosphorylation state of the shared response regulator (RR), DivK, thereby controlling asymmetric cell division.^8^ In *Vibrio harveyi*, quorum-sensing signals are integrated through multiple HKs that converge on LuxO, generating a ternary (three-state) output rather than a simple binary response.^11^ The information-processing capacity of these natural phosphosignaling architectures highlights their potential as blueprints for engineering advanced phosphosignaling logic circuits.

Despite the knowledge of these natural multikinase networks, synthetic TCS networks remain largely limited to single-input designs, with crosstalk minimized to preserve modularity. Three strategies have been explored to integrate signals based on TCSs: (1) a chimeric tandem sensor with two tandem sensory domains and a single HK module, integrating signals through conformational changes;^12^ (2) 2HK–2RR systems yielding either two distinct output proteins,^13^ a single split-protein output,^4^ or a common reporter under two distinct RR-regulated promoters;^14^ and (3) a 2HK–1RR system demonstrated in mammalian cells where two HKs converge on a shared RR, yet the presence of the cognate ligands was not necessary for phosphosignaling.^15^ These prior studies highlight the following critical gap: how can we engineer a bottom-up, modular approach for designing 2HK circuits to sense any signal of interest and support multi-input logic?

TCSs have been used to build single-input biosensors responsive to light^14, 16, 17^ and small molecules,^18–24^ providing a diverse parts toolkit for building two-input circuits. Here, we engineer a 2HK–1RR system (hereafter referred to as “2HK system” for brevity) in *E. coli* using simple TCS components, achieving programmable opto- and chemo-signal integration with graded, multi-level outputs. These synthetic networks recapitulate signal processing behaviors observed in natural systems, such as ternary output logic observed in quorum sensing 2HK circuits^11^, demonstrating that controlled crosstalk can be harnessed to build sophisticated decision-making systems in engineered cells.

## RESULTS

### Design of a two-histidine kinase phosphosignaling circuit with ternary output logic

To develop a modular system where two histidine kinases (HKs) phosphorylate a shared response regulator (RR), we co-expressed two copies of the cognate HK, each with a different N-terminal sensory domain responding to a distinct environmental signal. This protein engineering strategy, swapping the sensory domains of HKs, has been previously used to generate chimeric HKs regulated by new input signals.^17, 25, 26^ As a proof of concept, we constructed a 2HK system composed of the following two chimeric HKs: the blue-light sensor YF1 and the indole-3-aldehyde (I3A) sensor I3A-HK. Both chimeric HKs incorporate the same FixL-derived HK module and phosphorylate the same RR, the *Bradyrhizobium diazoefficiens* FixJ (Fig. 1a). The phosphorylated FixJ binds to the *fixK2* promoter to activate expression of the mCherry reporter gene (Fig. 1b). The output was characterized by mCherry fluorescence intensity normalized to the optical density at 660 nm, which is absent in the host strain background (Supplementary Fig. 1a). This design results in a 2HK phosphosignaling network that was expected to sense both blue light and I3A.

**Fig. 1.**
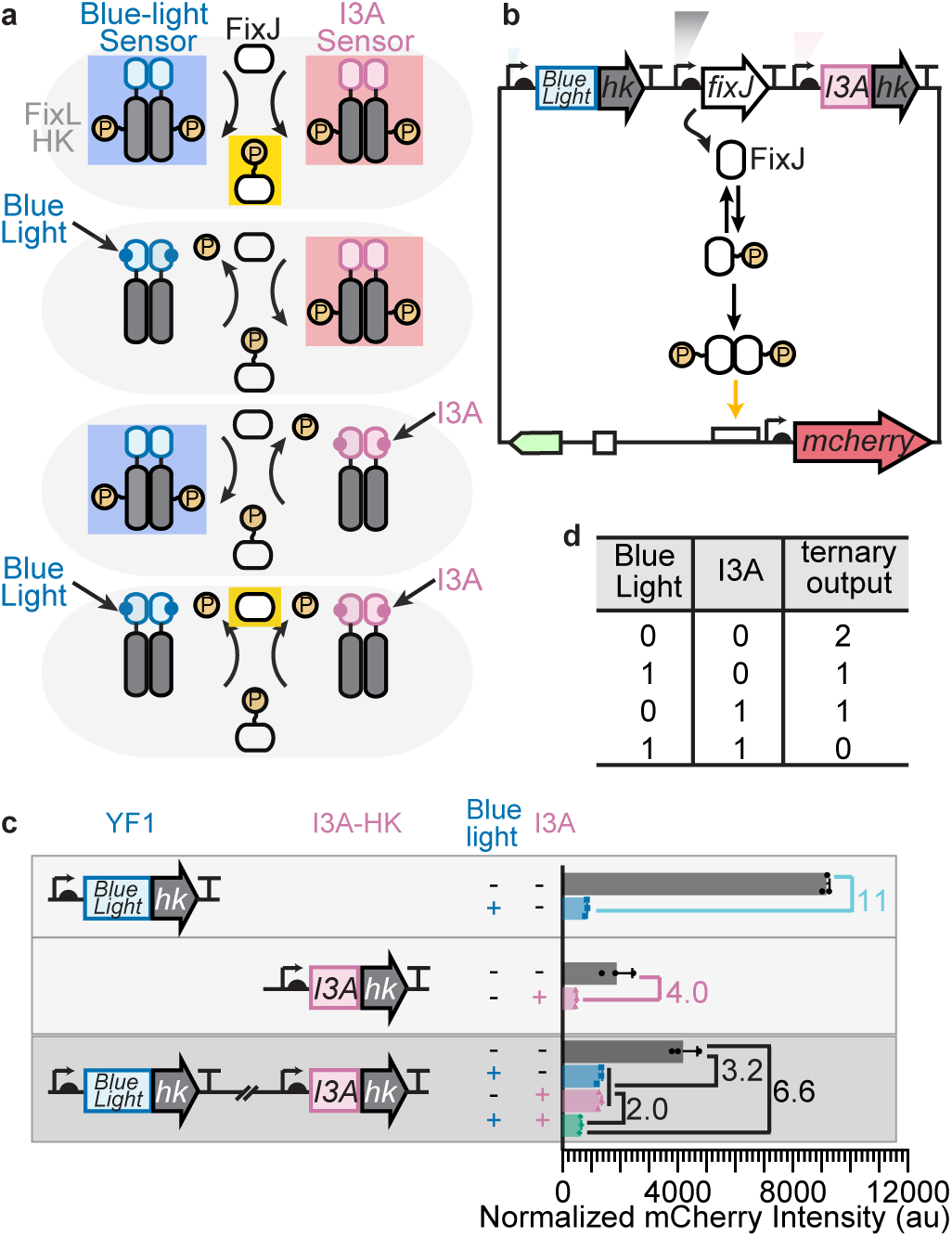
A phosphosignaling circuit composed of two histidine kinases (HKs) that phosphorylate a common response regulator (RR) processes blue-light and indole-3-aldehyde (I3A) signals with a ternary output response. a **Schematic of the 2HK–1RR** (hereafter “2HK”) system under four scenarios. Both HK modules of YF1 (blue-light sensor) and I3A-HK (I3A sensor) are derived from FixL, and both HKs form homodimers, regulating the phosphorylation of the shared RR FixJ. Each HK switches between a kinase state (shaded blue for YF1 and pink for I3A-HK) which phosphorylates FixJ; and a phosphatase state which dephosphorylates FixJ. Since both HKs are OFF-switches, when no input is present, RR is phosphorylated (yellow-shaded in the top scenario); when both inputs are present, RR is dephosphorylated (yellow-shaded in the bottom scenario). **b** Plasmid schematic for the 2HK system. mCherry expression is regulated by the dimerization state of FixJ, which depends on its phosphorylation. White rectangle: FixJ binding site; white square: origin of replication; green arrow: chloramphenicol resistance marker. **c** Normalized fluorescence (mCherry/OD_660_) for the YF1 blue light sensing 1HK–1RR (hereafter “1HK”) system (top), the I3A-HK 1HK system (middle) and the (YF1)-(I3A-HK) 2HK system (bottom) under four input conditions: no inputs (black), blue light only (blue), 100 µM I3A only (pink), and both inputs (green). In the 1HK systems, blue light caused 11-fold repression (*p* < 0.0001) and I3A caused a 4.0-fold repression (*p* ≈ 0.04). In the 2HK system, a single input caused 3.2-fold repression (*p* < 0.0001); a second input further reduced the output 2.0-fold (*p* ≈ 0.05); both inputs together led to 6.6-fold repression (*p* < 0.0001) relative to no-input. au: arbitrary unit. **d** Ternary output truth table for the 2HK system, mapping outputs to high (2), moderate (1) or low (0). Data represent mean ± standard deviation of three biological replicates.

To determine the appropriate expression levels of the HKs and the RR in the 2HK circuit, the plasmid was designed based on Golden Gate hierarchical assembly^27^ to enable efficient screening of promoters and ribosome binding sites (RBSs), with the backbone based on the pDusk vector architecture.^28^ As a starting point, both YF1 and I3A-HK were constitutively expressed from the SJM914 promoter and the PET RBS, and FixJ was constitutively expressed using the *lacI^Q^* promoter (hereafter referred to as “p1”) and a tailRBS that mimics its expression level in pDusk as closely as possible. However, this 2HK system responded only weakly to I3A in the presence of blue light (Supplementary Fig. 1b). Thus, we switched to the J23100 promoter for I3A-HK, which is reported to be stronger than SJM914 *in vitro*.^27^

We first verified the baseline activities of each 1HK–1RR system (hereafter referred to as “1HK system” for brevity) independently. Expression of the OFF-switch blue-light-sensing 1HK system (*yf1* gene) resulted in an 11-fold repression of the output when exposed to blue light (9126 ± 118 arbitrary units (au) vs 842 ± 36 au) (Fig. 1c, top). Further, constitutive expression of the OFF-switch I3A-HK yielded a 4.0-fold reduction in output when exposed to 100 µM I3A (1890 ± 542 au vs 474 ± 27 au) (Fig. 1c, middle).

We next evaluated the output from our engineered 2HK circuit with SJM914-PET RBS-YF1 and J23100-PET RBS-I3A-HK (Fig. 1c, bottom). With both inputs present (636 ± 30 au, green bar), the output decreased 6.6-fold compared to the no-input condition (4177 ± 497 au, black bar). Interestingly, each HK contributes equally, as the addition of either cognate input decreased the output 3.2-fold (4177 ± 497 au, black bar vs 1299 ± 91 au, blue bar, and 1296 ± 73 au, pink bar). In these single-input conditions, one HK acts in its kinase state, mediating phosphorylation of the RR, while the other acts in its phosphatase state, facilitating dephosphorylation of the RR, resulting in opposing effects on FixJ phosphorylation. Adding both inputs further decreased the output by another 2.0-fold relative to the single-input condition, establishing three distinct output states: high (no input), moderate (one input), and low (both inputs). This specific type of gate with a ternary output operates as an inverse count gate in which the output level is determined by the number of absent inputs (Fig. 1d), a behavior of particular interest for circuit design.^11, 29, 30^

Variation of the RR promoter in both 1HK (Supplementary Fig. 2a, b) and 2HK (Supplementary Fig. 2c, d) systems impacted the output response. Within the 2HK circuit, fold-change varied from 1-fold to 18-fold and output logic was quaternary (four-stage) or ternary. For downstream parameter analysis, we selected the p1-tailRBS promoter for *fixJ* as it was the only one that produced a ternary output. Thus, we concluded that both HK and RR promoters play a critical role in phosphosignaling circuit performance, and we chose the following promoter-RBS combinations for the three components: SJM914-PET for *yf1*, J23100-PET for *i3a-hk*, and p1-tailRBS for *fixJ*.

### Dimerization tags improve phosphosignaling output

In our 2HK system design, although the two HKs have distinct sensory domains with different dimerization interfaces, each HK contains the same dimerization and histidine-phosphotransfer (DHp) domain, allowing potential heterodimerization. Current understanding of the functional consequences of such heterodimers is limited,^31–33^ but could involve productive crosstalk for signal integration or inhibition of activity. Indeed, our 2HK system, compared to the blue-light sensing 1HK system with the absence of blue light (Fig. 1c top, black bar), exhibited a lower output (46%) when both inputs were absent (Fig. 1c bottom, black bar). This indicates that the co-expression of I3A-HK functionally interferes with YF1’s kinase activity.

To resolve this issue, we developed an approach to bias the system toward homodimer formation by fusing a high-affinity homodimerization tag with a dissociation constant substantially smaller than that of the DHp domain^34, 35^ to the N-terminus of the HKs (Fig. 2a). This design was intended to make homodimerization energetically favorable, thereby outcompeting DHp-mediated heterodimerization between the two different chimeric HKs. We utilized two distinct dimerization tags: 1) the GCN4 leucine zipper (LZ)^36, 37^ and 2) the *de novo* designed coiled-coil dimerization tag (CCDi), which utilizes a distinct dimerization interface^35^ (Supplementary Fig. 3a).^38^ A flexible glycine-serine linker (GGGGS) was introduced to prevent potential disruption of HK function observed with direct attachment.^39, 40^ In the 1HK control experiments, we observed that fusion of the LZ dimerization tag to the N-terminus of YF1 had little effect on the activity (Fig. 2b top vs. Fig. 1c top), whereas fusion of the CCDi dimerization tag to I3A-HK enhanced the output response (Fig. 2b middle vs. Fig. 1c middle).

**Fig. 2.**
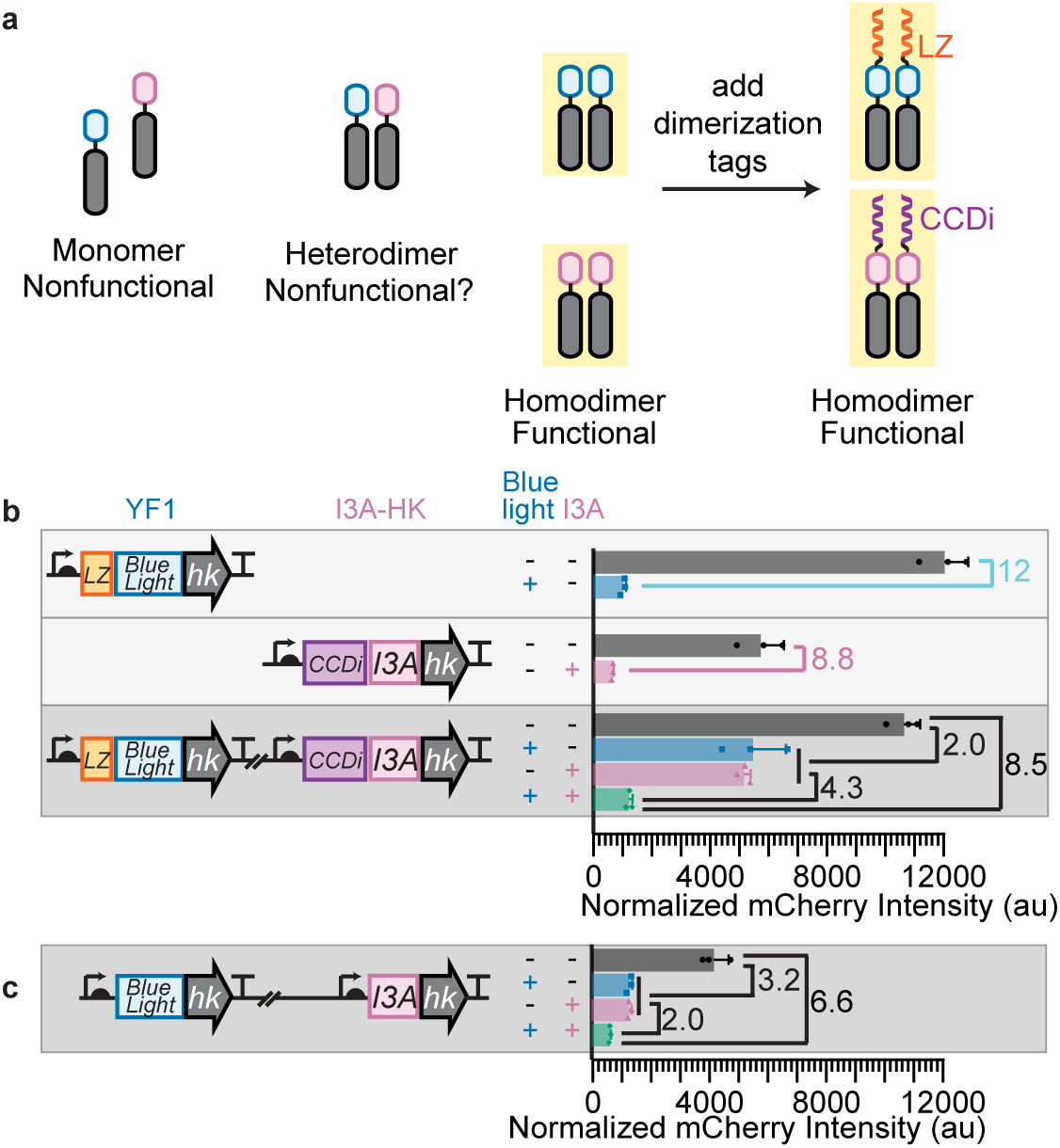
Dimerization tags reinforce homodimerization and enhance the dynamic range of the 2HK phosphosignaling circuit. **a** Schematic showing the oligomerization states of the HKs, which were hypothesized to be affected by the HK N-terminal dimerization tags. LZ: leucine zipper; CCDi: another coiled-coil dimerization tag. Blue: blue-light sensory domain; pink: I3A sensory domain; yellow shade: functional. **b** Effect of dimerization tags on normalized fluorescence (mCherry/OD_660_) for the 1HK LZ-YF1 system (top), the 1HK CCDi-I3A-HK system (middle), and the tagged 2HK system (bottom). In the 1HK systems, LZ-YF1 was repressed by blue light 12-fold (*p* = 0.0016) and CCDi-I3A-HK was repressed by I3A 8.8-fold (*p* = 0.0078). In the 2HK (LZ-YF1)-(CCDi-I3A-HK) system, single input caused 2.0-fold (*p* < 0.0001) repression, while the addition of a second input caused 4.3-fold (*p* < 0.001) repression; dual-input caused 8.5-fold repression (*p* < 0.0001) relative to no-input. **c** Normalized fluorescence (mCherry/OD_660_) of the tagless 2HK system from Fig. 1c (bottom) is shown for comparison. Data represent mean ± standard deviation of three biological replicates.

Introducing these two distinct dimerization tags into the 2HK system (Fig. 2b bottom) restored the output to levels comparable to the LZ-YF1 1HK system (Fig. 2b top) in the absence of the input (black bars), increasing the dynamic range of the 2HK system. This result suggests dimerization tags enhance the net kinase activity in the 2HK system, likely by reducing non-functional heterodimer formation. Interestingly, tagging only one HK was insufficient to restore the activity level of the corresponding 1HK systems (black bars of Supplementary Fig. 3b (LZ-YF1)-(I3A-HK) vs. Fig. 2b (LZ-YF1) and Fig. 1c (I3A-HK); Supplementary Fig. 3b (YF1)-(LZ-I3A-HK) vs. (YF1) and (LZ-I3A-HK)). These results suggest that both dimerization tags are needed to significantly reduce non-functional heterodimer formation, possibly due to steric clashes between LZ and CCDi that prevent heterodimer formation. Moreover, tagging both HKs with the same LZ tag, which would promote not only homodimerization but also heterodimerization, nearly abolished responsiveness to blue light (Supplementary Fig. 3b (LZ-YF1)-(LZ-I3A-HK)), indicating significant formation of nonfunctional heterodimers. In conclusion, having distinct dimerization tags on the two HKs enhanced the dynamic range in output.

A particularly informative condition for assessing potential heterodimer interference arises when only one input is present. This raises a structural question of how signal binding leads to histidine kinase autophosphorylation. When a signal binds to the sensory domain, it triggers a conformational change that is transmitted through the interdomain linker. This structural shift repositions the DHp and CA domains, thereby modulating HK activity.^41^ In a heterodimer, however, the two different monomers may impose opposing conformational changes on the linker between the sensory domain and the HK module, creating structural frustration within the HK module. Consistent with this model, when only one input is present, the output of the 2HK system with distinct dimerization tags (Fig. 2b bottom, blue bar and pink bar) is 4.1-fold higher than the system lacking dimerization tags (Fig. 2c, blue bar and pink bar).

Altogether, enforcing HK homodimerization via fusion of distinct dimerization tags expands the dynamic range of output and maintains a graded ternary response characteristic of an inverse count gate. With this established, we next examined how intrinsic biochemical properties of HKs shape signal integration in 2HK networks. Specifically, we tuned the kinase–phosphatase activity balance and introduced input-controlled ON-versus OFF-switches to investigate their influence on information flow.

### Tuning the phosphatase activity of the histidine kinases enables binary NAND logic

To program the logic behavior of the 2HK circuit, we first considered tuning the phosphatase activity of the HKs. Our mathematical modeling (see Methods section) suggests that when both HKs have a reduced phosphatase rate constant (Fig. 3a, yellow region), the repression of the output would require input stimulation of both HKs (Fig. 3a, green triangles; Supplementary Fig. 4a-b), producing a NAND gate response. This modulation switches the output logic from ternary to binary. Guided by this prediction, we designed and built 1HK and 2HK phosphosignaling circuits in which the HKs carry mutations that reduced phosphatase activity.

**Fig. 3.**
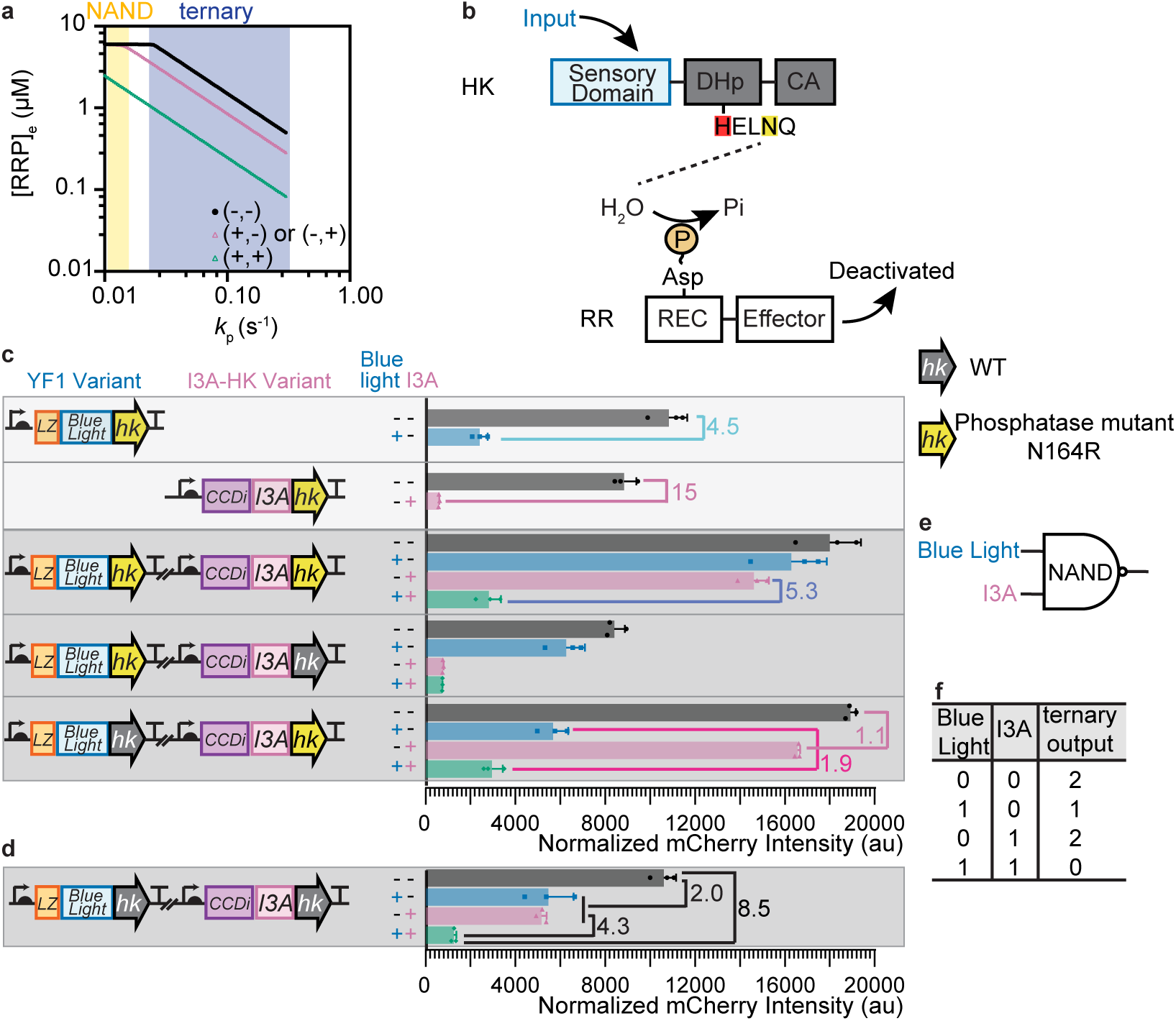
Dual phosphatase mutants alter 2HK signal integration to produce NAND logic. **a** Mathematical model simulation of 2HK systems (see Methods section). The y-axis shows steady state phosphorylated response regulator concentration ([RRP]_e_). At lower phosphatase activity (*k*_p_), the output logic transitions from ternary (blue region) toward binary logic (yellow region). Symbols indicate input conditions: black circle: both inputs absent; pink triangle: one input present; green triangle: both inputs present. **b** Schematic of the proposed mechanism of HK phosphatase activity. Residue N164 (yellow shaded) is near the active site histidine (red shaded). DHp: dimerization and histidine-phosphotransfer domain; CA: catalytic/ATP-binding subdomain; REC: receiver domain. **c** Normalized fluorescence (mCherry/OD_660_) showing effects of the N164R phosphatase mutation on 1HK or 2HK systems. LZ-YF1[N164R] 1HK system was repressed by blue light 4.5-fold (*p* = 0.0009); CCDi-I3A-HK[N164R] 1HK system was repressed by I3A 15-fold (*p* = 0.0014). The dual-N164R 2HK system displays repression only when both inputs are present (5.3-fold vs. “Dark, 100 μM I3A” condition; *p* < 0.0001). (LZ-YF1[N164R])-(CCDi-I3A-HK) had weak or no response to blue light (*p* = 0.0026 without I3A; *p* = 0.9999 with I3A), but responded strongly to I3A (*p* < 0.0001). (LZ-YF1)-(CCDi-I3A-HK[N164R]) weakly responded to I3A in dark conditions (1.1-fold; *p* = 0.0007), however responded to I3A more strongly in blue light conditions (1.9-fold; *p* = 0.0003). **d** Normalized fluorescence (mCherry/OD_660_) of the tagged wild type (WT) 2HK system from Fig. 2b for comparison. **e** NAND gate symbol for the dual-N164R system in Panel c. **f** Truth table for the (LZ-YF1)-(CCDi-I3A-HK[N164R]) system in Panel c. Data represent mean ± standard deviation of three biological replicates.

We targeted a conserved residue, N164, located near the phosphorylatable histidine in the HK module and implicated in coordinating a nucleophilic water in mediating RR dephosphorylation (Fig. 3b).^42–44^ Mutating residue N164 produced two distinct phenotypes.

Replacing it with an arginine (N164R) retained signal responsiveness in the 1HK systems but altered their activities. In YF1, N164R led to substantially increased output in the phosphatase state (blue bars in Fig. 3c top vs. Fig. 2b top) while the mutation significantly increased output in the kinase state for I3A-HK (black bars in Fig. 3c for CCDi-I3A-HK[N164R] vs. Fig. 2b middle). By contrast, mutating N164 to a glutamic acid (N164E) nearly abolished the phosphatase activity in both 1HK systems. YF1[N164E] exhibited constitutively high output and lost responsiveness to blue light (Supplementary Fig. 5 top). I3A-HK[N164E] had higher output compared to the WT, but with limited responsiveness (Supplementary Fig. 5 middle vs. Fig. 2b middle). These mutants thus provide tunable control points for circuit design.

We next incorporated the phosphatase mutants into both HKs in the 2HK system. In the dual-N164R system, we observed a binary NAND gate response (Fig. 3c). Specifically, when no or one input was present (black, blue and pink bars), the system maintained a high output. The substantial reduction of the output required the presence of both inputs (green bar) where both HKs were in the phosphatase state, which is different from the dual-WT 2HK circuit output (Fig. 3d). This conversion from ternary output to binary NAND gate output matched our model predictions (Fig. 3a). Meanwhile, when I3A-HK[N164E] is combined with YF1[N164R], this 2HK system displayed a leaky NAND gate (Supplementary Fig. 5, bottom), which is consistent with the high output and low fold-change of I3A-HK[N164E] (Supplementary Fig. 5, middle), underscoring the need to precisely reduce, but not eliminate, phosphatase activity to preserve the NAND logic.

Incorporation of the N164R phosphatase mutation into only one of the two HKs abolished NAND gate function (Fig. 3c), but also revealed additional design principles. YF1[N164R] in combination with a fully functional I3A-HK resulted in an output predominantly driven by I3A signaling. When I3A stimulates phosphatase activity in I3A-HK, the output remains low regardless of whether YF1 is in the kinase state (pink bar) or the phosphatase state (green bar). This suggests that YF1[N164R]’s kinase activity in its kinase state is insufficient to overcome the opposing phosphatase activity of I3A-HK, indicating that careful balancing of the kinase and phosphatase activities of both HKs is needed to ensure one HK does not dominate over the other.

In contrast, the combination of YF1 and I3A-HK[N164R] (Fig. 3c, bottom) exhibited another type of ternary output: high output in the dark (with or without I3A; black bar and pink bar), moderate output under blue light alone (blue bar), and low output when both inputs were present (green bar). This reflects a unique gating behavior in which I3A responsiveness is nearly abolished in the dark (1.1-fold change) but partially restored under blue light (1.9-fold change), demonstrating that modulating the activity of a single HK within a 2HK system can control signal flow, effectively programming the conditions under which I3A sensing is permitted or blocked. We describe this ternary output logic as a context “if [A=1], then [B MID OFF-switch]” gate (Supplementary Table 1), where repression by Input B (I3A) is only enabled in the presence of Input A (blue light). This reveals that 2HK architectures can be used to encode context-specific responses in synthetic circuits, through asymmetry in the activities of the two HKs. Together, these results indicate that precise tuning of phosphatase activity is essential for programming both binary and context-specific behaviors in 2HK circuits.

### Inverting histidine kinase responses expands circuit logic

To further broaden 2HK circuit capabilities, we converted the OFF-switch HKs to ON-switches, which activate output when their input is present (Fig. 4a). Strategies for reversing the response of the HK include introducing point mutations in the sensory domain^45^ and varying the length of the linker between the sensory domain and the HK module.^25, 46^ The former approach was applied to YF1, where an H22P mutation^45^ inverted its response, producing a 7.2-fold increase in the output upon blue-light stimulation (Fig. 4b, top). We next varied the linker length in I3A-HK, as this was more likely to be a generalizable approach to convert from OFF-switch to ON-switch HKs. We utilized primer-aided truncation of chimeric hybrids (PATCHY)^47^ to build a library of I3A sensors with various linker lengths (784 designs). One variant, I3A-6, which contains a 6-residue deletion in the linker, functioned as an ON-switch, exhibiting a 6.6-fold increase in output upon I3A stimulation (Fig. 4b).

**Fig. 4.**
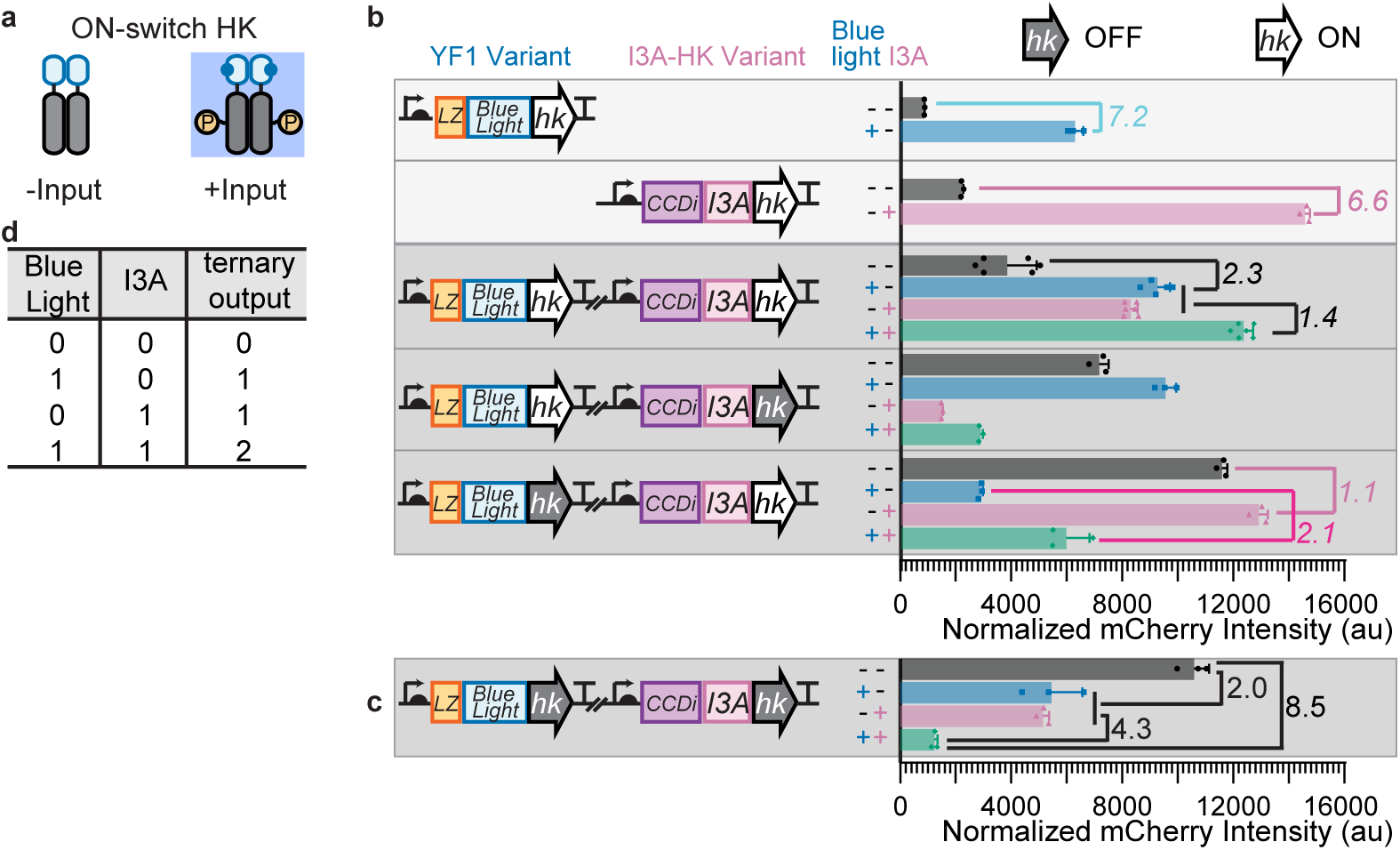
Dual ON-switches alter 2HK signal integration to produce a count gate. **a** Schematic of an ON-switch HK which is in the phosphatase state when the input is absent and switches to the kinase state when the input is present. **b** Normalized fluorescence (mCherry/OD_660_) showing the effect of ON-switch mutant on 1HK and 2HK output logic. ON-switch mutant YF1[H22P] 1HK system exhibited higher output under blue light (7.2-fold; *p* = 0.001). H22P is a point mutation at the C-terminal of the A’α helix at the N-terminal of YF1’s sensory domain, previously reported to confer ON switch behavior.^45^ The fold-change is italicized to highlight reversal relative to the OFF-switches. ON-switch mutant I3A-6 1HK system showed higher output in the presence of 100 μM I3A (6.6-fold; *p* < 0.0001). (I3A-6) contains a 6-amino-acid deletion in the linker between the sensory domain and the HK module relative to the original I3A sensor. The dual ON-switch 2HK system had higher output when blue light or I3A was present (3.0-fold; *p* < 0.0001); and the output was further increased when a second input was also present (1.4-fold; *p* < 0.0001). When YF1 ON-switch and I3A OFF-switch are paired together, the system had higher output when blue light was present (*p* < 0.0001 without I3A; *p* = 0.0008 with I3A). The YF1[OFF]-I3A[ON] system showed higher output when I3A was present. When blue light was absent, the change was not substantial (1.1-fold; *p* = 0.0313); when blue light was present, the change was substantial (2.1-fold; *p* = 0.0002). **c** Output of the tagged wildtype (WT) OFF-switch 2HK system from Fig. 2b for comparison. **d** Truth table for the dual ON-switch 2HK system, termed the “count gate”. HK module cartoons: gray fill = OFF-switch, white fill = ON-switch. Data represent mean ± standard deviation of three to six biological replicates. For the (LZ-YF1[H22P])-(CCDi-I3A-6) system, 6 colonies were analyzed except for the “blue-light, no I3A” condition, which had 5 due to one culture failing to grow. All other systems were measured in three replicates.

Based on these results, we incorporated ON-switch variants into the 2HK circuit and observed that this reversed the input–output polarity, producing activation rather than repression in response to input signals. Specifically, the dual ON-switch 2HK system produced the lowest output in the absence of blue light and I3A (black bar), while the addition of either input led to a 2.3-fold increase in output (blue bar and pink bar) (Fig. 4b), and the presence of both inputs (green bar) led to an additional 1.4-fold increase relative to the single-input scenario. This stepwise increase in output is the opposite of the dual OFF-switch system which leads to a stepwise decrease (Fig. 4c). We describe this additive response as a “count gate”, where the output reflects the number of inputs present (Fig. 4d).

Some natural 2HK systems combine ON- and OFF-switch HKs, as in the DivJ– PleC/DivK phosphosignaling circuit in *Caulobacter crescentus*, which drives asymmetric cell division through sequential checkpoint signals.^48^ Similarly, our synthetic circuit pairing an ON-switch HK and an OFF-switch HK produced quaternary or ternary output logics. The YF1[ON]+I3A[OFF] system (Fig. 4b) yielded four distinct outputs, so we named it “quaternary (four-stage) output logic”, although the circuit exhibited higher fold-change in response to I3A than to blue light. In comparison, the YF1[OFF]+I3A[ON] system produced a new type of ternary logic (Fig. 4b, bottom). In the absence of blue light, the system was unresponsive to I3A (black bar vs. pink bar). However, under blue light, I3A-induced activation was restored, resulting in a 2.1-fold increase in output (green bar vs. blue bar). This represents another example of a context gate, where Input B (I3A) is functionally gated by the environmental presence of Input A (blue light) (Supplementary Table 1). Access to these context gates typically arises from differences in the balance between kinase and phosphatase activity, suggesting that, beyond a change in signaling polarity, the I3A-HK ON-switch’s phosphatase activity is insufficient to counteract YF1’s kinase function. These results show that polarity inversion expands the range of achievable logic, enabling circuits with additive, conditional, or staged outputs.

## DISCUSSION

This work establishes a modular framework for programming signal integration via phosphosignaling circuits based on two-component systems. By engineering two-histidine kinase (2HK) systems that converge on a shared response regulator (RR), we identified several key tunable parameters that govern logic behaviors. These include the expression levels of the HKs and the RR, as well as each HK’s activity mode (kinase activity, phosphatase activity, ON-switch or OFF-switch). Tuning these features enabled circuit behaviors across a spectrum ranging from binary to ternary logic, including context-specific gating and additive “count gate” behaviors.

Our circuits recapitulate logic architectures observed in natural multikinase networks while also revealing novel modes of signal integration not previously described such as NAND gate behavior via phosphatase tuning and context-dependent sensing via asymmetric activity. For example, pairing two OFF-switch HKs produced a graded, ternary response similar to the *Vibrio harveyi* Lux system,^11^ in which two input signals must be integrated for full deactivation. In contrast, combining an ON-switch and an OFF-switch HK generated logic resembling the *Caulobacter crescentus* PleC–DivJ/DivK network,^8^ which is activated by one signal and repressed by the other, a configuration that drives asymmetric cell division in this organism. By opting for a bottom-up approach, we were successful in uncovering novel 2HK signaling modes that have never been reported, such as using phosphatase tuning to achieve context-dependent sensing and NAND gate behavior. In total, we built five (Figure 5a, 5b) of the 81 possible ternary output gates (Fig. 5c; Supplementary Table 1). Ternary outputs can replicate standard binary behavior (Supplementary Table 2) while also enabling 23 unique, context-dependent gates that implement IF–THEN logic. Twelve other possible gates either allow phosphosignaling circuits to count the number of active signals or detect specific transition, such as a shift from checkpoint-like Signal A to Signal B. These systems provide a foundation for layered logic in microbial circuits, where ternary outputs can reduce noise, enable sequential behaviors, or support analog-like responses.

**Fig. 5.**
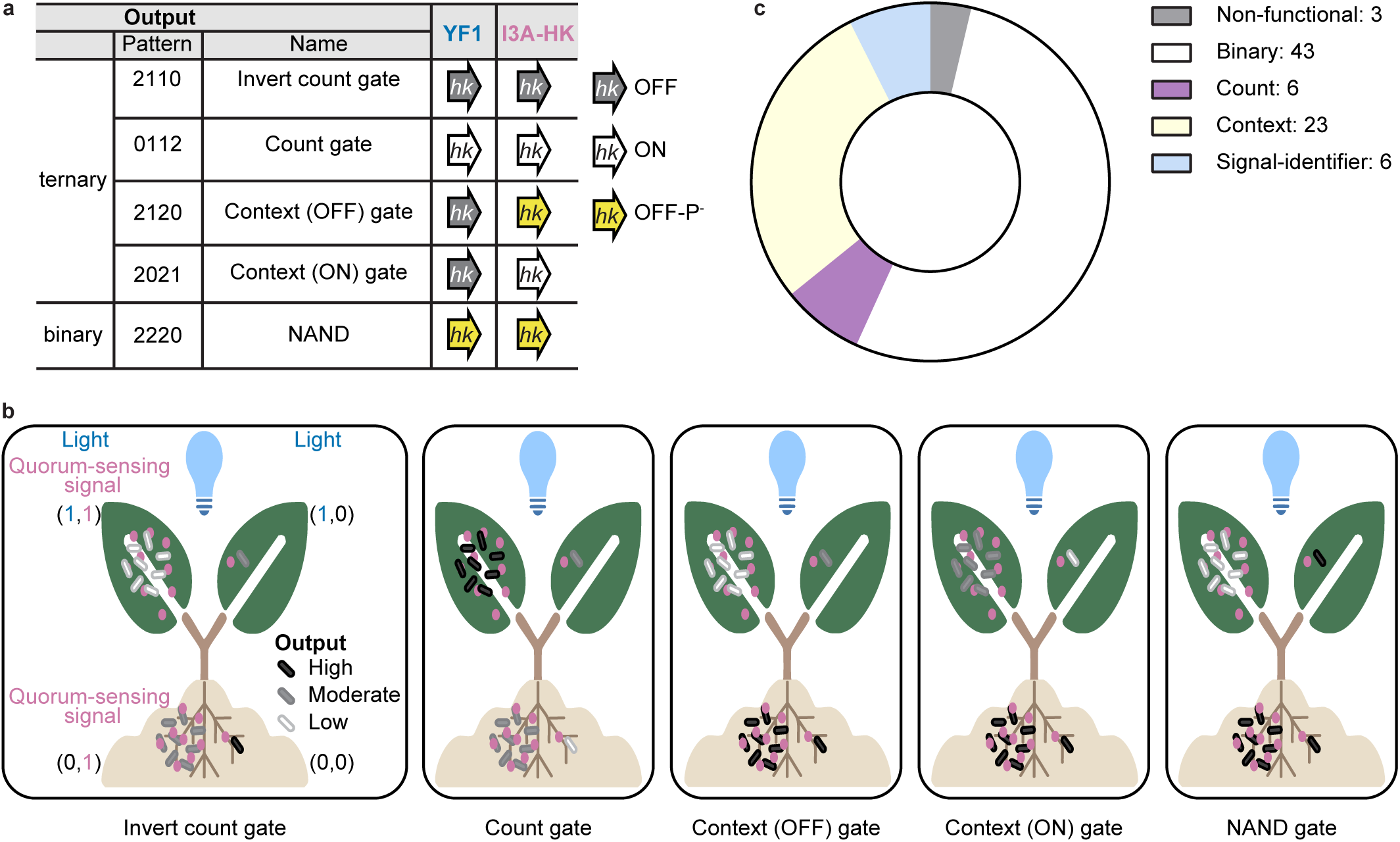
Programmability of bacterial phosphosignaling networks with ternary and binary logical outputs. **a** Achieved output logic behaviors of the 2HK blue light + I3A circuit in this work. Output patterns correspond to the four input states: (0,0), (1,0), (0,1) and (1,1). **b** Example application of the 2HK system to discriminate bacterial populations surrounding plants. Blue light distinguishes whether cells are inside or outside soil; indole or its derivatives report local cell density. The four environmental scenarios are: (0,0), bacteria is in soil and at low density; (1,0) outside soil and at low density; (0,1) in soil and at high density; (1,1) outside soil and at high density. Output levels are indicated by bacterial color: high (black), moderate (gray), low (white). **c** Distribution of the five categories of ternary logic gates (see Supplementary Table 1).

Several challenges and opportunities remain before these phosphosignaling systems can be broadly applied. First, while HKs encode thousands of distinct sensors, we only know the ligand binding specificity and ON/OFF switch behavior of a small subset of these sensors, which limits the capacity to track signals of interest. Second, achieving balanced outputs often requires tuning multiple parameters (e.g., HK activity and system promoters), which can be labor-intensive. In particular, when designing chimeric HKs, we lack a clear understanding of how chimeric junctions alter kinase/phosphatase activities, which remains an opportunity area for further research.^25, 46, 49–51^ Finally, most synthetic TCS platforms remain focused on transcriptional outputs and are not yet adapted for rapid, post-translational control of the target, constraining their ability to mediate time-sensitive responses. Even so, the advantages of our 2HK system are significant. Compared to transcription factor-based logic, which operates at the timescales and energy consumption of gene expression of the components, 2HK systems offer fast, ATP-driven post-translational control. When layered with transcriptional circuits, they provide a route to multi-tiered signal processing and sophisticated cellular programming.

These capabilities create opportunities across synthetic biology. For example, ternary logic could regulate the expression of two distinct genes in a way that, at low FixJ concentrations, only the high-affinity promoter binds to FixJ; while at high FixJ concentrations, both high- and low-affinity promoters are activated. A circuit integrating light and indole cues could distinguish above-ground plant-associated bacteria from dense soil communities, activating outputs only in the intended population, an approach valuable for biosensing and biocontainment in dynamic environments (Fig. 5b). Multi-stage logic could also enable programmable biocontainment, where early warning cues trigger low-burden responses such as growth arrest, while stronger signals escalate to irreversible cell lysis. Similar staged architectures could support programmable probiotics that tune immune activity in steps, or metabolic circuits that coordinate sequential biosynthetic pathways. They might even guide the formation of engineered biofilms with spatial patterning based on unique spatial cues. Overall, our results show that multi-stage signaling architectures are feasible and reliable, offering routes to advanced decision-making in engineered microbes for applications in biosensing, biocontainment, probiotics, metabolic engineering and biofilm patterning.

## METHODS

### Molecular cloning

#### Golden Gate Assembly

The 1HK and 2HK plasmids were constructed using a modular Golden Gate assembly-based system (Supplementary Tables 3 and 4) by using the EcoFlex kit which was a gift from Paul Freemont (Addgene kit #1000000080).^27^ Our customized version has three defined plasmid levels: Level 0 holds standardized BioParts such as promoters, RBSs, HK ORFs, RR ORFs, dimerization tags, and terminators; Level 1 combines these BioParts into HK or RR transcription units (TUs); and Level 2 integrates multiple TUs into complete 1HK or 2HK gene circuits.

YF1 and I3A-HK share identical amino acid sequences in their HK modules. Using identical DNA sequences risks homologous recombination. To mitigate this issue, we recoded the DNA sequence of the HK module in the Level 0 BioPart vectors to reduce sequence identity.

#### Unwanted restriction enzyme digestion sites were mutated out

The Level 0 part vectors were either constructed by other cloning methods (shown below), or ordered from Twist Bioscience (see Supplementary Table 5). The Level 1 destination vectors (DV) were entirely from the EcoFlex kit.^27^ The Level 2 destination vectors (DV) were completely customized.

The Golden Gate assembly reactions were done by using the NEBridge Golden Gate Assembly Kit BsaI-HF v2 (NEB, USA) (for Level 1 assembly) or the NEBridge Golden Gate Assembly Kit BsmBI-v2 (for Level 2 assembly). More specifically, the reaction mixture contains 50 ng of the destination vector and 100 ng of each part vector. The mixture of the components was incubated in a thermal cycler. The setting of the thermal cycler was as follows: for Level 1 assembly, there were 30 cycles of cut (37 °C, 1 min or 5 min) and ligate (16 °C, 1 min or 5 min), followed by deactivation of the enzyme (60 °C, 5 min); for Level 2 assembly, the program is similar, only replacing 37 °C with 42 °C. We found 5-min incubation usually resulted in a higher probability of introducing transposons into the plasmid, so all subsequent reactions were performed with 1-min incubation time.

#### Gibson Assembly

The backbone and the insert were linearized by PCR, and the length of the product was confirmed by running an agarose gel, followed by either PCR Cleanup or gel extraction. Then the template was digested by DpnI. For the Gibson assembly reaction, the master mix was combined with 0.07-0.08 pmol vector and the insert-to-vector molar ratio was 1:5. Water was added to make the total volume around 20 µL. The mixture was incubated at 50 °C for 1 hour, followed by keeping on ice for 10 min. Detailed information is in Supplementary Table 5-8.

#### Splicing by Overlap Extension (SOE)

The backbone is linearized by PCR, and the insert is either embedded in the primers or amplified from a template. Then the two fragments were assembled by overlap-extension PCR.

#### NEBridging

Briefly, a single-stranded DNA (ordered from IDT) is used to “bridge” the two ends of a linearized backbone. The kit (NEBuilder® HiFi DNA Assembly Master Mix) was purchased from NEB (catalog number E2621S). 1 µL of 100 µM single-stranded DNA was mixed with 499 µL TE buffer, resulting in 0.2 µM oligo. The reaction mixture contained 5 µL 0.2 µM oligo, 1 µL 0.005 pmol/µL linearized backbone, 4 µL water, and 10 µL 2X NEBuilder HiFi DNA assembly master mix. The mixture was incubated at 50 °C for 1 hour, followed by transformation into competent cells. The backbone was linearized by PCR and purified by either gel extraction (Qiagen, USA) or PCR Cleanup (Qiagen, USA).

#### QuikChange

The Agilent QuikChange II kit was used. Briefly, the reaction mixture contained 50 ng template, 125 ng forward primer and 125 ng reverse primer, besides the buffer, dNTP and water to make the total volume 50 µL. Then 2.5 U PfuUltra HF DNA polymerase was added, followed by the reaction which included the initial denaturation (95 °C, 30 s) and the thermal cycling (Denaturation at 95 °C for 30 s; annealing at 55 °C for 60 s; extension at 68 °C for 60 s/kb; 18 cycles for multiple amino acid deletion or insertion). Then the reaction product was incubated on ice for 2 min, and 10 U DpnI was added, followed by centrifugation for 1 min and incubation at 37 °C for 1 hour. The digestion product was transformed into Top10 chemically competent cells. pDusk was a gift from Andreas Möglich (Addgene plasmid # 43795; http://n2t.net/addgene:43795; RRID:Addgene_43795)

### Bacterial Strains and Growth

1. *E. coli* plasmids, primers, and strains used in this work are listed in Supplementary Tables 3, 5, 8, and 9. Primers were designed using j5, SnapGene, or PrimerX and purchased from Integrated DNA Technologies (IDT, Coralville, IA). Gene blocks were synthesized by Twist Bioscience (South San Francisco, CA). Customized Biopart Vectors for Golden Gate assembly were constructed by either the authors or Twist Bioscience. Site-directed mutations were generated by QuikChange. Plasmids were purified by the Qiagen miniprep kit (Qiagen, USA).
2. *E. coli* strains were grown in LB media or on LB-agar plates (1.5% w/v agar). Antibiotic or other chemical working concentrations: ampicillin: 50 ng/μL for liquid media, 100 ng/μL for agar plates; kanamycin: 30 ng/μL for liquid media; 50 ng/μL for agar plates; chloramphenicol: 20 ng/μL for liquid media; 30 ng/μL for agar plates. IPTG for Golden Gate assembly plates: 0.1 mM; X-gal for Golden Gate assembly plate: 40 ng/μL. Cultures were done in 10 mL culturing tubes.

### Light Apparatus

Blue light was applied from an LED (470 nm; purchased from DigiKey; Catalog LXML-PB01-0040) located at the top of a small incubator. The plug was made by the University of Pittsburgh Electronics Shop.

### Chemicals

Indole-3-aldehyde (I3A) used in this study was purchased from Thermo Scientific Chemicals and stored at 4 °C. The 100 mM solution was prepared with DMSO as the solvent, filtered through a 0.2 μm-pore nylon sterile syringe filter, and stored at −20 °C.

### Fluorescence Reporter Assay

Plasmids were transformed into chemically competent cells of *E. coli* Top10. Colonies were obtained by plating the transformed cells onto selective LB-agar plates or by streaking from glycerol freezer stocks of the strain onto selective LB-agar plates, and growing overnight at 37 °C. Three single colonies were picked and inoculated into 200 µL LB media in 1.7 mL

Eppendorf tubes with appropriate antibiotics, and these seed cultures were incubated at 37 °C with 275 rpm shaking for around 10 hours. Then 50 µL seed culture was added to 100 µL LB media in a clear 96-well microplate (Greiner Bio-One, Catalog number 655101), and then the optical density at 660 nm (OD_660_) was measured by a Tecan M1000 plate reader and normalized to 0.01 by adding an appropriate volume into LB media with appropriate antibiotics. (We chose 660 nm instead of 600 nm because mCherry has significant absorption at 600 nm, which would interfere with the measurement of cell optical density.) The normalized culture was then aliquoted to the culturing tubes which were put under testing conditions. The cultures were incubated at 37 °C, 275 rpm. After 12.5 hours, 200 µL culture was transferred from the culturing tube to the well of a black-bottom, black-wall 96-well microplate (Greiner Bio-One, Catalog number 655076). Then 50 µL culture was transferred from the well of the black microplate to the well of a clear 96-well microplate which already contained 100 µL LB in it. Then OD_660_ (by the clear microplate) and red fluorescence (by the black microplate; excitation wavelength: 585 ± 5 nm; emission wavelength: 610 ± 5 nm; Gain: 120) were measured by Tecan M1000 plate reader.

### Statistical Analysis

The mean and the standard deviation of normalized mCherry intensity were calculated by Microsoft Excel and graphs were generated by GraphPad Prism 10 (GraphPad Software, San Diego, USA). Fold-changes of the constructs are listed in Supplementary Table 10.

All data are from at least three biological replicates. The comparisons between two groups were done by Welch’s t-test. The comparisons involving four groups were done by ordinary one-way ANOVA with Tukey’s multiple comparisons test. Statistical analyses were done in GraphPad Prism 10. Important *p* values are reported in the figure legends.

### Generation of an I3A-HK ON-switch

The PATCHY (primer-aided truncation of chimera hybrids) protocol^46, 47^ was utilized to identify an I3A-HK variant with an inverted response to indole-3-aldehyde. The protocol was performed on (pBL1), which was modified from pXYSq10 (Supplementary Table 5) to contain both linkers from the parental HKs for the sensory domain and the HK module. The PATCHY library was purified using the DNA Clean & Concentrator-5 kit (ZymoResearch #D4013) and eluted in 10 µL; all of which was transformed into 25 µL of *E. cloni* 10G SUPREME Electrocompetent cells (Biosearch Technologies) using manufacturer’s recommendations. The library was amplified by inoculating the transformation reaction into 50 mL of LB-kanamycin (50 µg/mL) liquid media, grown at 30 °C for 24 hours at 250 rpm in a 250 mL baffled flask. The naïve library was isolated using a ZymoResearch MIDIprep kit from the pelleted culture, and >98% of the potential 784 (28 × 28) variants in the theoretical library were detected by next-generation sequencing (SeqCenter). Electrocompetent *E. coli* BW25993 (ref 52) (30 µL; obtained from the *E. coli* Genetic Resource Center, Cheshire, CT, USA) was transformed with 2 µL of the PATCHY library and plated on LB-agar plates containing kanamycin and I3A at a final concentration of 50 µg/µL and 100 µM, respectively. Transformants were grown at 30 °C for 48 hours, followed by incubation at 4 °C for 48 hours to allow for mCherry fluorescence development. Colonies were picked and resuspended in 100 µL of 1X phosphate buffered saline (PBS), followed by inoculating 25 µL of resuspended culture into 1 mL of LB-kanamycin or LB-kanamycin containing 100 µM I3A in a deep 96-well plate. The cultures were grown at 30 °C, shaking at 800 rpm. After 24 hours, cultures were diluted 10-fold into 200 µL of 1X PBS, where OD600 and red fluorescence were measured as above. Of the 384 colonies picked, 21 colonies exhibited a >10-fold increase in red fluorescence in the presence of I3A versus without I3A. Sanger sequencing of the linker region from the top 3 clones revealed them to have the same linker sequence (DITARLQELQSELVHVSRLSA), which we designate I3A-6.

### Mathematical Modeling

COPASI (COmplex PAthway SImulator) was used to simulate the reactions. Briefly, the species and the reactions were input into the software. The parameters (concentrations and rate constants)^53, 54^ are listed in Supplementary Table 11. Then the steady-state was simulated. The resulting RRP concentration of each scenario was recorded. We examined a scenario in which the activities and concentrations of both HKs were identical, using default parameters from Supplementary Table 11.

Two rate constants—autophosphorylation rate constant (*k_a_ ^+^*) and dephosphorylation rate constant (*k*_p_)—are particularly important. We considered the scenario where only the autophosphorylation rate constant (*k_a_ ^+^*) changes between kinase and phosphatase states, while other parameters, including *k*_p_, remain constant. Higher *k_a_ ^+^*in the kinase state underlies the increased kinase activity in this configuration.

## DATA AVAILABILITY

Data are available from the corresponding author upon request.

## ACKNOWLEDGEMENTS

This work was supported by the U.S. Department of Energy, Office of Science, Office of Biological and Environmental Research, Lawrence Livermore National Laboratory (LLNL) BioSecure SFA within the Secure Biosystems Design program. Work at LLNL is performed under the auspices of the U.S. Department of Energy at Lawrence Livermore National Laboratory under Contract DE-AC52-07NA27344. Shangri-La Hou received 2025 Wass Family Summer Research Fellowship Award from the University of Pittsburgh. We thank Dr. Dante Ricci for early efforts in the project conception and project advice. We thank Dr. Andreas Möglich’s lab at the University of Bayreuth for developing the blue-light sensor YF1 and their variants, the pDusk plasmid, and for developing the PATCHY cloning method. We thank Dr. Paul S. Freemont’s lab at Imperial College London and Addgene for the EcoFlex MoClo kit (Addgene Kit # 1000000080). We thank Dr. Alex Deiters’ lab at the University of Pittsburgh for sharing the Tecan M1000 plate reader.

## AUTHOR CONTRIBUTIONS

W.S.C. and X.S. conceived the research and analyzed the data; W.S.C. and Y.J. acquired funding and supervised the study. X.S. designed, constructed and characterized the plasmids and performed the related data analysis, and performed the mathematical modeling. B.L. constructed and characterized the ON-switch version of the I3A sensor. S.H. helped with cloning and characterization. X.S., W.S.C. and B.L. wrote the manuscript. Y.J., M.C.Y. and S.H. edited the manuscript.

## COMPETING INTERESTS

The authors declare no competing financial interests.

**Correspondence** and requests for materials should be addressed to W. Seth Childers.

