## Supplementary Information for "Programmable multiplexed phosphosignaling networks in bacteria"

Table of Contents

Supplementary Figure 1 - Normalized mCherry intensity of the no-plasmid strain and the dual-SJM914 promoter strain.

Supplementary Figure 2 - Modulating RR expression levels reshaped how the 1HK or 2HK systems integrate and process blue light and I3A signals.

Supplementary Figure 3 - Distinct LZ and CCDi dimerization tags are needed for each HK to retain full dynamic range and responsiveness to both inputs.

Supplementary Figure 4 - Mathematical modeling of the 2HK system and parameter scan.

Supplementary Figure 5 - Evaluation of the N164E phosphatase variant mutation on the I3A-HK.

Supplementary Table 1 - All 81 Possible logic functions of a two-input, ternary-output circuit.

Supplementary Table 2 - All 16 Possible logic functions of a two-input, binary-output circuit.

Supplementary Table 3 - Plasmids for characterization used in this study.

Supplementary Table 4 - Level 2 Golden Gate plasmid assembly methods.

Supplementary Table 5 - Plasmids that are not for characterization used in this study.

Supplementary Table 6 - Plasmid assembly detailed descriptions.

Supplementary Table 7 - Gene blocks used in this study.

Supplementary Table 8 - DNA oligos used in this study.

Supplementary Table 9 - *E. coli* strains used in this study.

Supplementary Table 10 - Fold-changes for the circuits.

Supplementary Table 11 - Default mathematical modeling parameters for the 2HK-1RR system.

Supplementary References

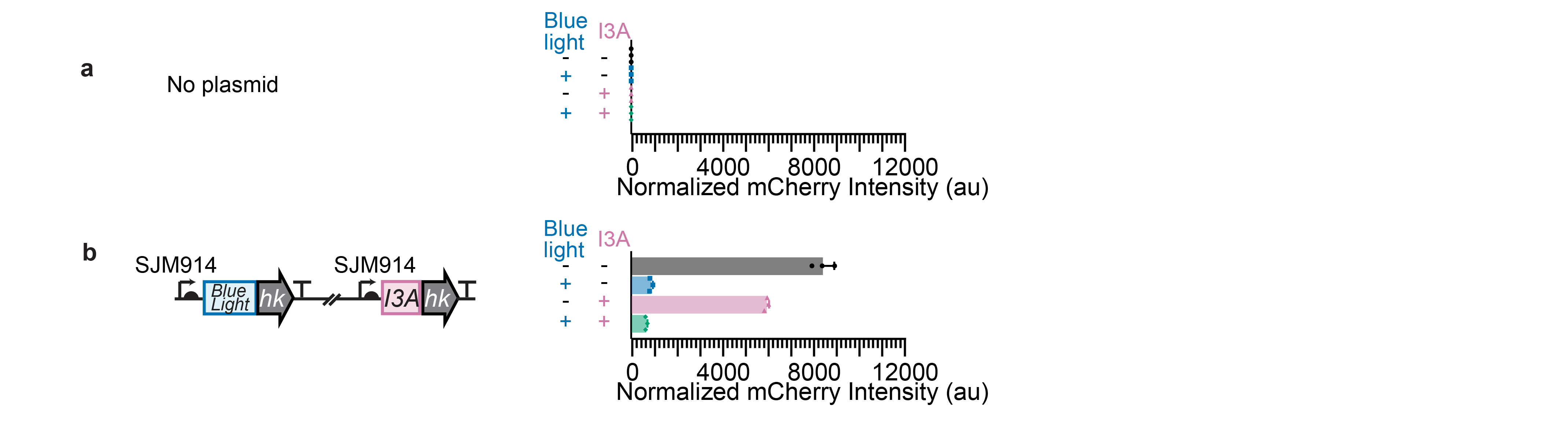

**Supplementary Figure 1. Normalized mCherry intensity of the no-plasmid strain and the dual-SJM914 promoter strain. a** *E. coli* Top10 lacking plasmids. **b** The 2HK circuit in which both the *yf1* gene and *i3a-hk* gene (without dimerization tags) are expressed constitutively from the SJM914 promoter with a PET ribosome binding site. The *fixJ* response regulator gene is expressed from the p1 promoter with the tailRBS ribosome binding site. In dark, I3A addition produced only a 1.4-fold repression (*p* < 0.0001) relative to no I3A; and in blue light, I3A addition yielded a 1.3-fold change (not significant). Data represent the mean ± standard deviation of three biological replicates.

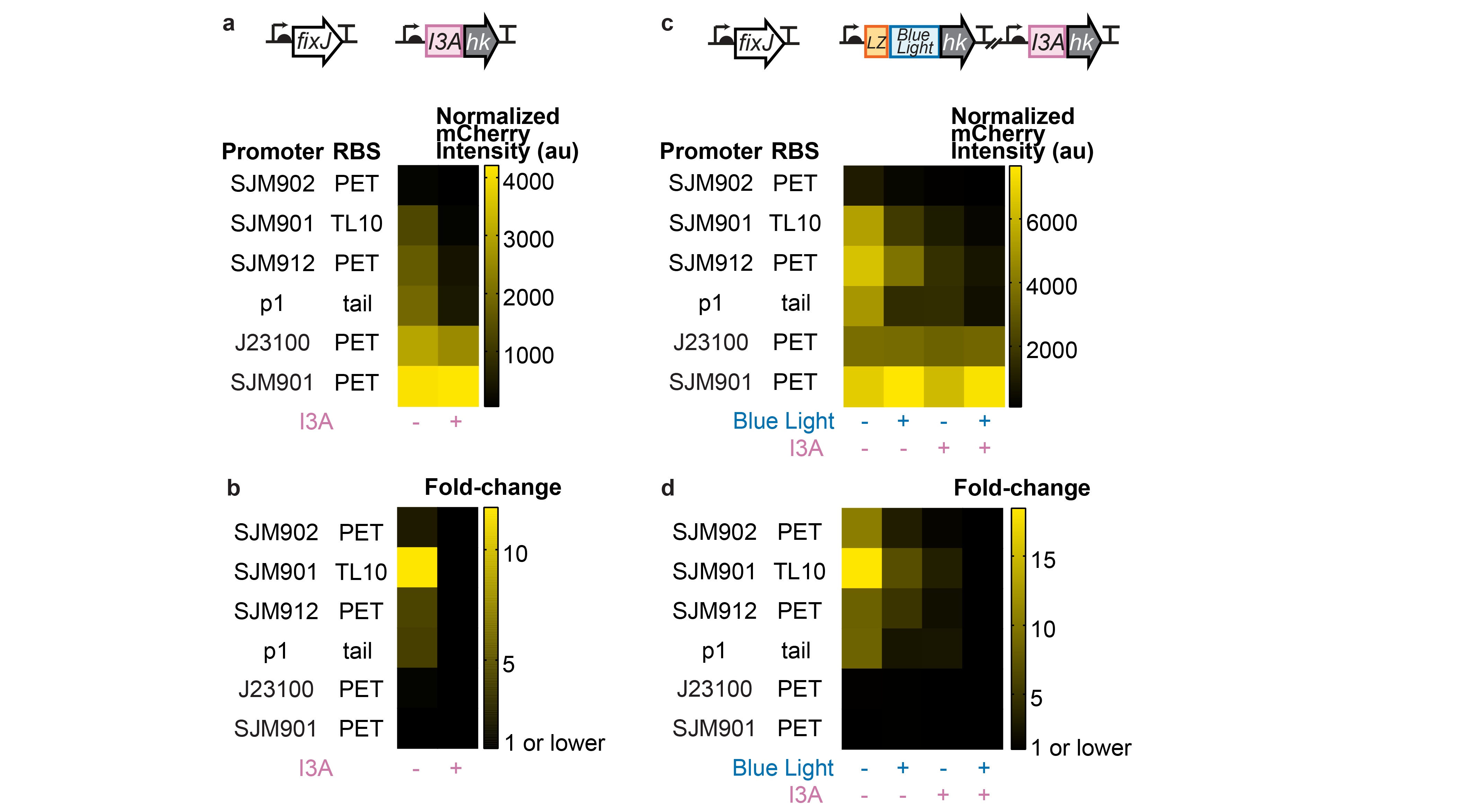

**Supplementary Figure 2. Modulating RR expression levels reshaped how the 1HK or 2HK systems integrate and process inputs.** (**a**) Normalized fluorescence intensity (mCherry/OD_660_) and (**b**) fold-change for J23100-PET-I3A-HK 1HK systems in *E. coli* Top10 with varied promoters and ribosome binding sites for the *fixJ* RR gene. The fold-change (output without I3A / output with I3A) was the highest at 11.9-fold (*p* = 0.0174) when using SJM901-TL10. When cellular FixJ concentration is too high, FixJ dimerizes in the absence of phosphorylation, resulting in gene activation both with and without a signal. p1-tailRBS data are reused from Fig. 1c, processed here as a heat map for comparison. (**c**) Normalized fluorescence intensity (mCherry/OD_660_) and (**d**) fold-change (output under a given condition / output in the presence of both inputs) for (SJM914-PET-LZ-YF1)-(J23100-PET-I3A-HK) 2HK systems in *E. coli* Top10 using the same set of varied promoters and ribosome binding sites as the 1HK system in (a) for *fixJ*. Data represent the mean of three biological replicates.

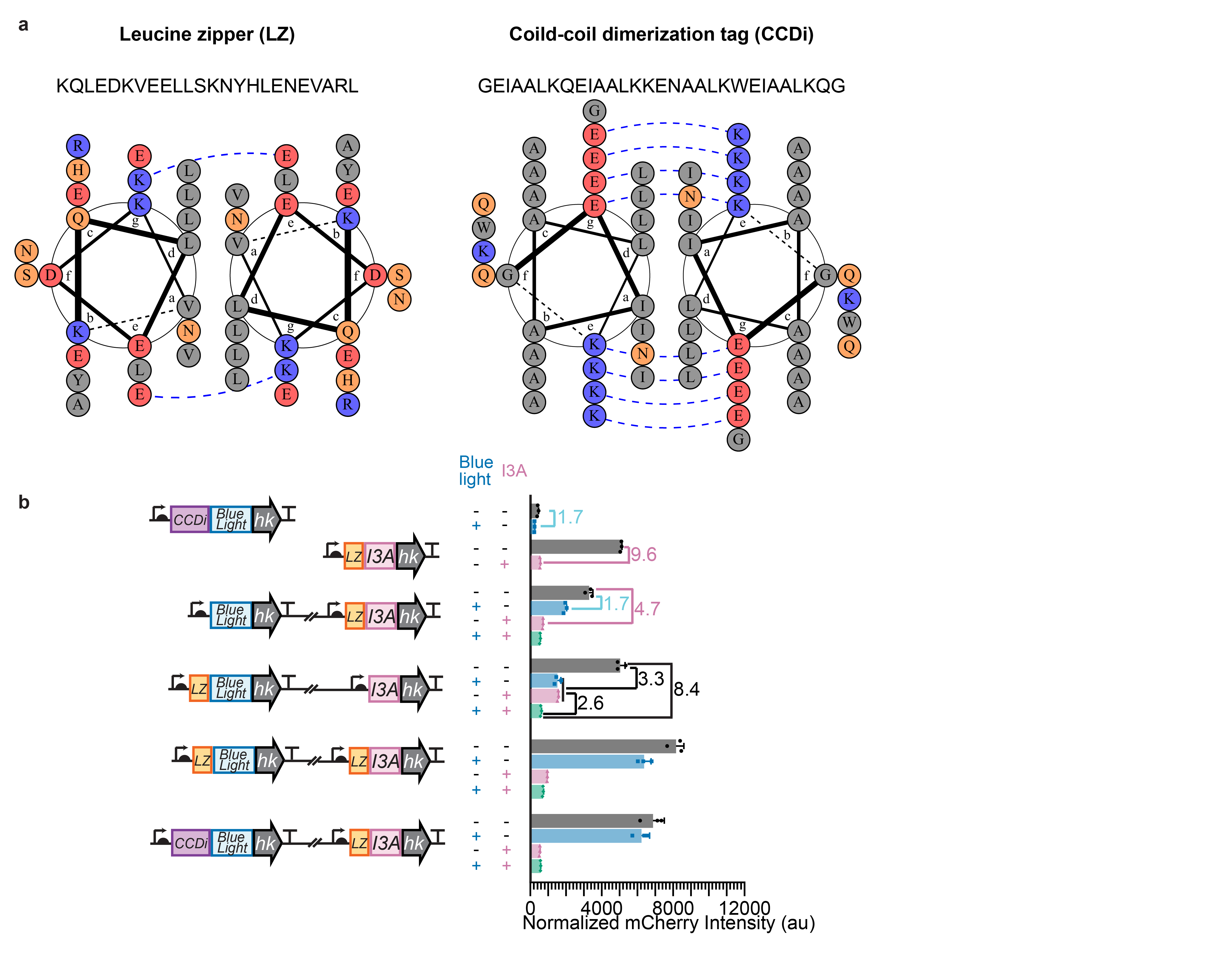

**Supplementary Figure 3.** **Distinct LZ and CCDi dimerization tags are needed for each HK to retain full dynamic range and responsiveness to both inputs. a** Heptad wheel of LZ dimer (left) and CCDi dimer (right) drawn using DrawCoil 1.0 software developed by Keating et al^1^. **b** Normalized fluorescence intensity (mCherry/OD_660_) for the CCDi-YF1 1HK system, LZ-I3A-HK 1HK system, (YF1)-(LZ-I3A-HK) 2HK system, (LZ-YF1)-(I3A-HK) 2HK system, (LZ-YF1)-(LZ-I3A-HK) 2HK system and (CCDi-YF1)-(LZ-I3A-HK) 2HK system. (LZ-YF1)-(I3A-HK) data are reused from Supplementary Fig. 2c, processed here as a bar graph for comparison. Data represent the mean ± standard deviation of three biological replicates.

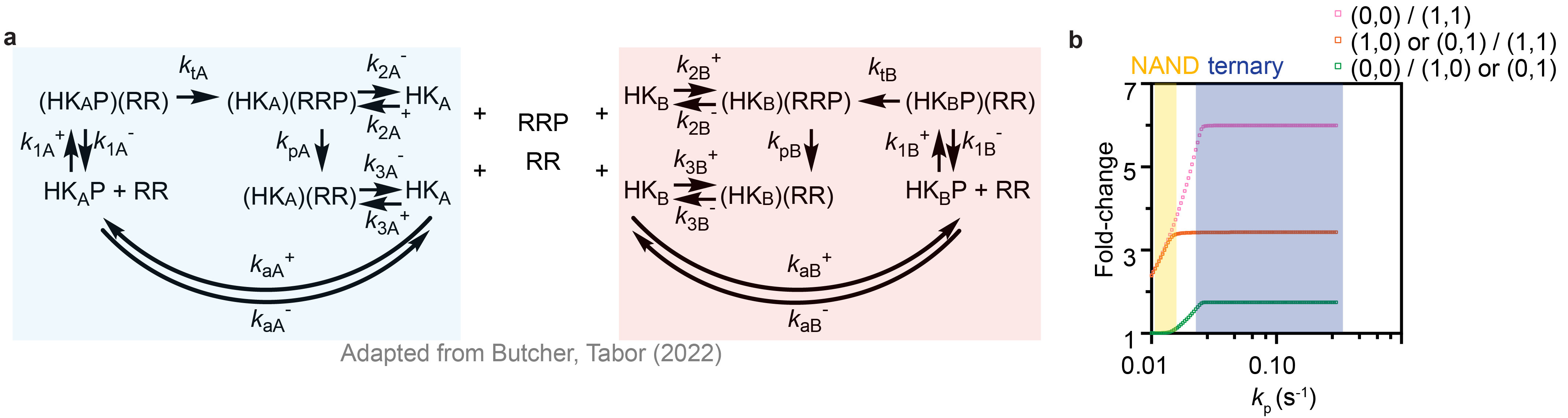

**Supplementary Figure 4. Mathematical modeling of the 2HK system and parameter scan. a** Reaction diagrams illustrating the modeled reactions of the two HKs (HK_A_: blue; HK_B_: pink), adapted from Butcher and Tabor (2022).^2^ Parameter definitions are in Supplementary Table 11. **b** Parameter scan of the phosphatase rate constant *k*_p_. y-axis: fold-changes. Both HKs are OFF-switches. Pink square: the output at (0,0) (no input present) over the output at (1,1) (both inputs present); orange square: the output at (1,0) or (0,1) over the output at (1,1); green square: the output at (0,0) over the output at (1,0) or (0,1). Blue shade: at this range of *k*_p_, the output is ternary; yellow shade: at this range of *k*_p_, the output is binary NAND gate.

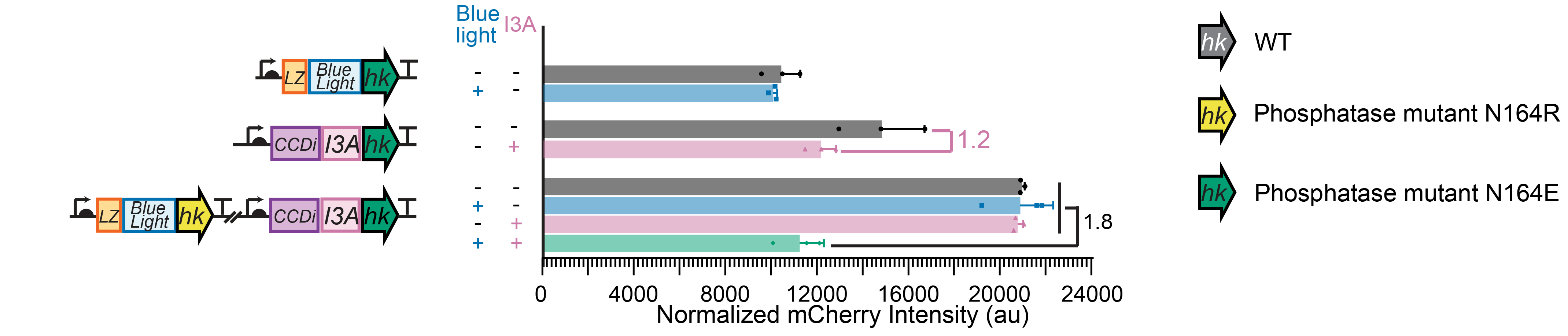

**Supplementary Figure 5. Evaluation of the N164E phosphatase mutation.** Normalized fluorescence intensity (mCherry/OD_660_) for the LZ-YF1[N164E] 1HK system, the CCDi-I3A-HK[N164E] 1HK system, and the (LZ-YF1[N164R])-(CCDi-I3A-HK[N164E]) 2HK system. The (LZ-YF1[N164R])-(CCDi-I3A-HK[N164E]) 2HK system displays a mild 1.8-fold repression (*p* < 0.0001) when both signals are present. Data represent the mean ± standard deviation of three biological replicates.

**Supplementary Table 1. All 81 possible logic functions of a two-input, ternary-output circuit.**

Binary input order: (A,B) = (0,0), (1,0), (0,1), (1,1). Each input condition yields a ternary output of 0, 1, or 2.

Ternary outputs fall into five categories: non-functional, binary, context, count, and graded binary gates.

1) Non-functional gates remain always ON or always OFF.
2) Binary: Some ternary gates produce binary-like patterns indistinguishable from true binary gates.
3) Count gate: When not binary & Output [(1,0)] = Output [(0,1)] & Output [(0,0)] ≠ Output [(1,1)].
4) Context gates follow the nomenclature “if [Condition], then [Effect]”.
5) Signal identifier gates detect a switch from signal A to signal B (encoded as 0, 1, or 2) with the third output state indicating “all” or “none.”

| **#** | **Gate Category** | **Description** | **Ternary Output Pattern*** |
| --- | --- | --- | --- |
| 0 | Non-functional | FALSE (Always 0) | 0000 |
| 1 | Binary | MID AND | 0001 |
| 2 | Binary | HIGH AND | 0002 |
| 3 | Binary | MID A AND NOT B | 0100 |
| 4 | Binary | MID BUFFER A | 0101 |
| 5 | Context | if [A=1], then [B LEAKY ON-switch] | 0102 |
| 6 | Binary | HIGH A AND NOT B | 0200 |
| 7 | Context | if [A=1], then [B LEAKY OFF-switch] | 0201 |
| 8 | Binary | HIGH BUFFER A | 0202 |
| 9 | Binary | MID NOT A AND B | 0010 |
| 10 | Binary | MID BUFFER B | 0011 |
| 11 | Context | if [B=1], then [A LEAKY ON-switch] | 0012 |
| 12 | Binary | MID XOR | 0110 |
| 13 | Binary | MID OR | 0111 |
| 14 | Count Gate | Count | 0112 |
| 15 | Signal Identifier | A only=2, B only=1 | 0210 |
| 16 | Context Gate | if [B=0], then [A HIGH ON-switch] | 0211 |
| 17 | Context Gate | if [A=0], then [B MID ON-switch] | 0212 |
| 18 | Binary | HIGH NOT A AND B | 0020 |
| 19 | Context Gate | if [B=1], then [A MID OFF-switch] | 0021 |
| 20 | Binary | HIGH BUFFER B | 0022 |
| 21 | Signal Identifier | A only=1, B only=2 | 0120 |
| 22 | Context Gate | if [A=0], then [B HIGH ON-switch] | 0121 |
| 23 | Context Gate | if [B=0], then [A MID ON-switch] | 0122 |
| 24 | Binary | HIGH XOR | 0220 |
| 25 | Count Gate | [0, 2, 1] Count Gate  (1 input high=2, 2 inputs=1, 0 inputs=0) | 0221 |
| 26 | Binary | HIGH OR | 0222 |
| 27 | Binary | MID NOR | 1000 |
| 28 | Binary | MID XNOR | 1001 |
| 29 | Count Gate | [1,0,2] Count Gate  (2 input high=2, 1 inputs=0, 2 inputs=2) | 1002 |
| 30 | Binary | MID NOT B | 1100 |
| 31 | Binary | MID IMPLY (B->A) | 1101 |
| 32 | Context Gate | if [B=1], then [A HIGH ON-switch] | 1102 |
| 33 | Context Gate | if [B=0], then [A LEAKY ON-switch] | 1200 |
| 34 | Signal Identifier | A only=2, B only=0 | 1201 |
| 35 | Context Gate | if [A=0], then [B MID OFF-switch] | 1202 |
| 36 | Binary | MID NOT A | 1010 |
| 37 | Binary | MID IMPLY (A->B) | 1011 |
| 38 | Context Gate | if [A=1], then [B HIGH ON-switch] | 1012 |
| 39 | Binary | MID NAND | 1110 |
| 40 | Non-functional | MID TRUE | 1111 |
| 41 | Binary | LEAKY AND | 1112 |
| 42 | Context Gate | if [A=1], then [B HIGH OFF-switch] | 1210 |
| 43 | Binary | LEAKY A AND NOT B | 1211 |
| 44 | Binary | LEAKY BUFFER A | 1212 |
| 45 | Context Gate | if [A=0], then [B LEAKY ON-switch] | 1020 |
| 46 | Signal Identifier | A only=0, B only=2 | 1021 |
| 47 | Context Gate | if [B=0], then [A MID OFF-switch] | 1022 |
| 48 | Context Gate | if [B=1], then [A HIGH OFF-switch] | 1120 |
| 49 | Binary | LEAKY NOT A AND B | 1121 |
| 50 | Binary | LEAKY BUFFER B | 1122 |
| 51 | Count Gate | [1,2,0] Count Gate  (0 input=2, 2 input=1, 1 inputs=0) | 1220 |
| 52 | Binary | LEAKY XOR | 1221 |
| 53 | Binary | LEAKY OR | 1222 |
| 54 | Binary | HIGH NOR | 2000 |
| 55 | Count Gate | [2,0,1] Count Gate  (1 input=2, 0 input=1, 2 inputs=0) | 2001 |
| 56 | Binary | HIGH XNOR | 2002 |
| 57 | Context Gate | if [B=0], then [A LEAKY OFF-switch] | 2100 |
| 58 | Context Gate | if [A=0], then [B HIGH OFF-switch] | 2101 |
| 59 | Signal Identifier | A only=1, B only=0 | 2102 |
| 60 | Binary | HIGH NOT B | 2200 |
| 61 | Context Gate | if [B=1], then [A MID ON-switch] | 2201 |
| 62 | Binary | HIGH IMPLY (B->A) | 2202 |
| 63 | Context Gate | if [A=0], then [B LEAKY OFF-switch] | 2010 |
| 64 | Context Gate | if [B=0], then [A HIGH OFF-switch] | 2011 |
| 65 | Signal Identifier | A only=0, B only=1 | 2012 |
| 66 | Count Gate | Inverse Count | 2110 |
| 67 | Binary | LEAKY NOR | 2111 |
| 68 | Binary | LEAKY XNOR | 2112 |
| 69 | Context Gate | if [B=1], then [A MID OFF-switch] | 2210 |
| 70 | Binary | LEAKY NOT B | 2211 |
| 71 | Binary | LEAKY IMPLY (B->A) | 2212 |
| 72 | Binary | HIGH NOT A | 2020 |
| 73 | Context Gate | if [A=1], then [B MID ON-switch] | 2021 |
| 74 | Binary | HIGH IMPLY (A->B) | 2022 |
| 75 | Context Gate | if [A=1], then [B MID OFF-switch] | 2120 |
| 76 | Binary | LEAKY NOT A | 2121 |
| 77 | Binary | LEAKY IMPLY (A->B) | 2122 |
| 78 | Binary | HIGH NAND | 2220 |
| 79 | Binary | LEAKY NAND | 2221 |
| 80 | Non-functional | Non-functional - HIGH TRUE | 2222 |

*: Binary Input (A,B) = (0,0), (1,0), (0,1), (1,1).

**Supplementary Table 2. All 16 possible logic functions of a two-input, binary-output circuit.**

| **#** | **Name** | **Output Pattern*** |
| --- | --- | --- |
| 0 | Constant FALSE | 0000 |
| 1 | AND | 0001 |
| 2 | A AND NOT B | 0100 |
| 3 | BUFFER A | 0101 |
| 4 | NOT A AND B | 0010 |
| 5 | BUFFER B | 0011 |
| 6 | XOR | 0110 |
| 7 | OR | 0111 |
| 8 | NOR | 1000 |
| 9 | XNOR | 1001 |
| 10 | NOT B | 1100 |
| 11 | IMPLY (B->A) | 1101 |
| 12 | NOT A | 1010 |
| 13 | IMPLY (A->B) | 1011 |
| 14 | NAND | 1110 |
| 15 | Constant TRUE | 1111 |

*: Order of input (A,B) scenarios: (0,0); (1,0); (0,1); (1,1).

**Supplementary Table 3. Plasmids for characterization used in this study.**

| **Plasmid ID** | **YF1** | | | **I3A-HK** | | | **FixJ promoter**–**RBS** |
| --- | --- | --- | --- | --- | --- | --- | --- |
|  | **promoter**–**RBS** | **tag** | **mutation** | **promoter**–**RBS** | **tag** | **mutation** |  |
| pXYSG2-33 | SJM914-PET | None | WT | NA | NA | NA | p1-tailRBS |
| pXYSG2-37 | NA | NA | NA | J23100-PET | None | WT | p1-tailRBS |
| pXYSG2-72 | SJM914-PET | None | WT | J23100-PET | None | WT | p1-tailRBS |
| pXYSG2-68 | SJM914-PET | LZ | WT | NA | NA | NA | p1-tailRBS |
| pXYSG2-132 | NA | NA | NA | J23100-PET | CCDi | WT | p1-tailRBS |
| pXYSG2-136 | SJM914-PET | LZ | WT | J23100-PET | CCDi | WT | p1-tailRBS |
| pXYSG2-137 | SJM914-PET | LZ | N164R | NA | NA | NA | p1-tailRBS |
| pXYSG2-195 | NA | NA | NA | J23100-PET | CCDi | N164R | p1-tailRBS |
| pXYSG2-189 | SJM914-PET | LZ | N164R | J23100-PET | CCDi | N164R | p1-tailRBS |
| pXYSG2-196 | SJM914-PET | LZ | N164R | J23100-PET | CCDi | WT | p1-tailRBS |
| pXYSG2-193 | SJM914-PET | LZ | WT | J23100-PET | CCDi | N164R | p1-tailRBS |
| pXYSG2-114 | SJM914-PET | LZ | H22P | NA | NA | NA | p1-tailRBS |
| pXYSG2-190 | NA | NA | NA | J23100-PET | CCDi | I3A-6 | p1-tailRBS |
| pXYSG2-183 | SJM914-PET | LZ | H22P | J23100-PET | CCDi | I3A-6 | p1-tailRBS |
| pXYSG2-181 | SJM914-PET | LZ | H22P | J23100-PET | CCDi | WT | p1-tailRBS |
| pXYSG2-182 | SJM914-PET | LZ | WT | J23100-PET | CCDi | I3A-6 | p1-tailRBS |
| pXYSG2-198 | SJM914-PET | NA | WT | SJM914-PET | None | WT | p1-tailRBS |
| pXYSG2-111 | NA | NA | NA | J23100-PET | None | WT | SJM902-PET |
| pXYSG2-108 | NA | NA | NA | J23100-PET | None | WT | SJM901-TL10 |
| pXYSG2-70 | SJM914-PET | LZ | WT | J23100-PET | None | WT | p1-tailRBS |
| pXYSG2-112 | NA | NA | NA | J23100-PET | None | WT | J23108-PET |
| pXYSG2-174 | NA | NA | NA | J23100-PET | None | WT | J23100-PET |
| pXYSG2-76 | NA | NA | NA | J23100-PET | None | WT | SJM901-PET |
| pXYSG2-150 | SJM914-PET | LZ | WT | J23100-PET | None | WT | SJM902-PET |
| pXYSG2-128 | SJM914-PET | LZ | WT | J23100-PET | None | WT | SJM901-TL10 |
| pXYSG2-149 | SJM914-PET | LZ | WT | J23100-PET | None | WT | SJM912-PET |
| pXYSG2-151 | SJM914-PET | LZ | WT | J23100-PET | None | WT | J23108-PET |
| pXYSG2-159 | SJM914-PET | LZ | WT | J23100-PET | None | WT | J23100-PET |
| pXYSG2-147 | SJM914-PET | LZ | WT | J23100-PET | None | WT | SJM901-PET |
| pXYSG2-197 | SJM914-PET | CCDi | WT | NA | NA | NA | p1-tailRBS |
| pXYSG2-67 | NA | NA | NA | J23100-PET | LZ | WT | p1-tailRBS |
| pXYSG2-89 | SJM914-PET | None | WT | J23100-PET | LZ | WT | p1-tailRBS |
| pXYSG2-173 | SJM914-PET | LZ | WT | J23100-PET | LZ | WT | p1-tailRBS |
| pXYSG2-172 | SJM914-PET | CCDi | WT | J23100-PET | LZ | WT | p1-tailRBS |
| pXYSG2-194 | NA | NA | NA | J23100-PET | CCDi | N164E | p1-tailRBS |
| pXYSG2-187 | SJM914-PET | LZ | N164R | J23100-PET | CCDi | N164E | p1-tailRBS |
| pXYSG2-192 | SJM914-PET | LZ | WT | J23100-PET | CCDi | N164E | p1-tailRBS |

**Supplementary Table 4. Level 2 Golden Gate plasmid assembly methods.**

| **Plasmid ID** | **Destination Vector** | **Part Vectors** | | |
| --- | --- | --- | --- | --- |
|  |  | **for YF1** | **for FixJ** | **for I3A-HK** |
| pXYSG2-32 | pXYSj32 | pXYSG1-55 | pXYSG1-11 | NA |
| pXYSG2-37 | pXYSqj43 | NA | pXYSG1-11 | pXYSG1-92 |
| pXYSG2-67 | pXYSqj43 | pXYSG1-97 | pXYSG1-11 | NA |
| pXYSG2-68 | pXYSj32 | pXYSG1-98-b | pXYSG1-11 | NA |
| pXYSG2-70 | pXYSj31 | pXYSG1-98-b | pXYSG1-11 | pXYSG1-92 |
| pXYSG2-72 | pXYSj31 | pXYSG1-80 | pXYSG1-11 | pXYSG1-92 |
| pXYSG2-76 | pXYSqj43 | NA | pG1-92 | pG1-120 |
| pXYSG2-83 | pXYSj32 | pXYSG1-104 | pXYSG1-11 | NA |
| pXYSG2-84 | pXYSj32 | pXYSG1-105 | pXYSG1-11 | NA |
| pXYSG2-89 | pXYSj31 | pXYSG1-80 | pXYSG1-11 | pXYSG1-97 |
| pXYSG2-108 | pXYSqj43 | NA | pG1-92 | pG1-135 |
| pXYSG2-111 | pXYSqj43 | NA | pG1-92 | pG1-138 |
| pXYSG2-112 | pXYSqj43 | NA | pG1-92 | pG1-139 |
| pXYSG2-114 | pXYSj32 | pXYSG1-129 | pXYSG1-11 | NA |
| pXYSG2-128 | pXYSj31 | pXYSG1-98-b | pXYSG1-135 | pXYSG1-92 |
| pXYSG2-132 | pXYSqj43 | NA | pXYSG1-11 | pG1-146 |
| pXYSG2-136 | pXYSj31 | pXYSG1-98-b | pXYSG1-11 | pG1-146 |
| pXYSG2-137 | pXYSj32 | pXYSG1-131 | pXYSG1-11 | NA |
| pXYSG2-147 | pXYSj31 | pXYSG1-98-b | pXYSG1-120 | pXYSG1-92 |
| pXYSG2-149 | pXYSj31 | pXYSG1-98-b | pXYSG1-137 | pXYSG1-92 |
| pXYSG2-150 | pXYSj31 | pXYSG1-98-b | pXYSG1-138 | pXYSG1-92 |
| pXYSG2-151 | pXYSj31 | pXYSG1-98-b | pXYSG1-139 | pXYSG1-92 |
| pXYSG2-159 | pXYSj31 | pXYSG1-98-b | pXYSG1-156 | pXYSG1-92 |
| pXYSG2-172 | pXYSj31 | pXYSG1-151 | pXYSG1-11 | pG1-97 |
| pXYSG2-173 | pXYSj31 | pXYSG1-98-b | pXYSG1-11 | pG1-97 |
| pXYSG2-174 | pXYSqj43 | NA | pG1-92 | pG1-156 |
| pXYSG2-175 | pXYSj32 | pXYSG1-160 | pXYSG1-11 | NA |
| pXYSG2-181 | pXYSj31 | pXYSG1-129 | pXYSG1-11 | pXYSG1-146 |
| pXYSG2-182 | pXYSj31 | pXYSG1-98-b | pXYSG1-11 | pXYSG1-148 |
| pXYSG2-183 | pXYSj31 | pXYSG1-129 | pXYSG1-11 | pXYSG1-148 |
| pXYSG2-187 | pXYSj31 | pXYSG1-131 | pXYSG1-11 | pXYSG1-165 |
| pXYSG2-189 | pXYSj31 | pXYSG1-131 | pXYSG1-11 | pXYSG1-166 |
| pXYSG2-190 | pXYSqj43 | NA | pXYSG1-11 | pXYSG1-148 |
| pXYSG2-192 | pXYSj31 | pXYSG1-98-b | pXYSG1-11 | pXYSG1-165 |
| pXYSG2-193 | pXYSj31 | pXYSG1-98-b | pXYSG1-11 | pXYSG1-167 |
| pXYSG2-194 | pXYSqj43 | NA | pXYSG1-11 | pXYSG1-165 |
| pXYSG2-195 | pXYSqj43 | NA | pXYSG1-11 | pXYSG1-167 |
| pXYSG2-196 | pXYSj31 | pXYSG1-131 | pXYSG1-11 | pXYSG1-146 |
| pXYSG2-197 | pXYSj32 | pXYSG1-151 | pXYSG1-11 | NA |
| pXYSG2-198 | pXYSj31 | pXYSG1-80 | pXYSG1-11 | pXYSG1-171 |

**Supplementary Table 5. Plasmids used in this study (not for characterization).**

| **Category** | **Plasmid** | **Description** | **Cloning Method / Reference** | **Addgene Plasmid #** |
| --- | --- | --- | --- | --- |
| Entry Vector |  | pBP-lacZ | Freemont (2016)* | 72948 |
|  |  | pBP-ORF |  | 72949 |
| Part Vector – Promoter |  | pBP-SJM914 |  | 72975 |
|  |  | pBP-J23100 |  | 72963 |
|  |  | pBP-SJM901 |  | 72966 |
|  |  | pBP-SJM912 |  | 72974 |
|  |  | pBP-SJM902 |  | 72967 |
|  |  | pBP-SJM915 |  | 72976 |
|  |  | pBP-SJM910 |  | 72972 |
| Part Vector – RBS |  | pBP-PET |  | 72981 |
|  |  | pBP-TL8 |  | 72992 |
|  |  | pBP-TL10 |  | 72994 |
| Part Vector – promoter + RBS | pXYSj11 | pBP-p1 | SOE | NA |
| Part Vector – tag | pXYSj15 | pBP-LZ | SOE | NA |
|  | pXYST78 | pBP-CCDi | Twist | NA |
| Part Vector – ORF | pXYSj23 | pBP-tailRBSFixJ | Gibson | NA |
|  | pXYST23 | pBP-YF1 | Twist | NA |
|  | pXYST55 | pBP-I3A | Twist | NA |
|  | pXYST24 | pBP-YF1[H22P] | Twist | NA |
|  | pXYST28 | pBP-YF1[N164R] | Twist | NA |
|  | pXYST48 | pBP-I3A[N164R] | Twist | NA |
|  | pXYST47 | pBP-I3A[N164E] | Twist | NA |
| Part Vector – terminator |  | pBP-L3S2P21 | Freemont (2016) | 72999 |
|  |  | pBP-L2U2H09 |  | 73007 |
|  |  | pBP-BBa_B0015 |  | 72998 |
| Level 1 Destination Vector |  | pTU1-A-lacZ |  | 72935 |
|  |  | pTU1-B-lacZ |  | 72936 |
|  |  | pTU1-C-lacZ |  | 72937 |
| Level 2 Destination Vector | pXYSj32 | pDusk2-a-RFP | GA | NA |
|  | pXYSj31 | pDusk2-b-RFP | GA | NA |
|  | pXYSqj43 | pDusk2-c-RFP | QuikChange | NA |
| Level 1 Construct | pXYSG1-80 | pTU1-A-(SJM914-PET-YF1) | GGA | NA |
|  | pXYSG1-98-b | pTU1-A-(SJM914-PET-LZ-YF1) |  |  |
|  | pXYSG1-131 | pTU1-A-(SJM914-PET-LZ-YF1[N164R]) |  |  |
|  | pXYSG1-129 | pTU1-A-(SJM914-PET-LZ-YF1[H22P]) |  |  |
|  | pXYSG1-151 | pTU1-A-(SJM914-PET-CCDi-YF1) |  |  |
|  | pXYSG1-92 | pTU1-C-(J23100-PET-I3A) |  |  |
|  | pXYSG1-146 | pTU1-C-(J23100-PET-CCDi-I3A) |  |  |
|  | pXYSG1-167 | pTU1-C-(J23100-PET-CCDi-I3A[N164R]) |  |  |
|  | pXYSG1-148 | pTU1-C-(J23100-PET-CCDi-I3A-6) |  |  |
|  | pXYSG1-171 | pTU1-C-(SJM914-PET-I3A) |  |  |
|  | pXYSG1-97 | pTU1-C-(J23100-PET-LZ-I3A) |  |  |
|  | pXYSG1-165 | pTU1-C-(J23100-PET-CCDi-I3A[N164E]) |  |  |
|  | pXYSG1-11 | pTU1-B-(p1-tailRBSFixJ) |  |  |
|  | pXYSG1-138 | pTU1-B-(SJM902-PET-FixJ) |  |  |
|  | pXYSG1-135 | pTU1-B-(SJM901-TL10-FixJ) |  |  |
|  | pXYSG1-137 | pTU1-B-(SJM912-PET-FixJ) |  |  |
|  | pXYSG1-139 | pTU1-B-(J23108-PET-FixJ) |  |  |
|  | pXYSG1-156 | pTU1-B-(J23100-PET-FixJ) |  |  |
|  | pXYSG1-120 | pTU1-B-(SJM901-PET-FixJ) |  |  |
| Only for cloning | pDusk | pDusk | Möglich (2012)** | 43795 |
|  | pXYSq2 | pDusk with mCherry | QuikChange | NA |
|  | pXYSq26-C2 | pXYSq2[c1373a][g2843a] | QuikChange | NA |
|  | pXYSq27-B | pXYSq2 without BsaI or BsmBI sites | QuikChange | NA |
|  | pXYSq10 | pDusk with mCherry and p3 promoter | GA | NA |
| PATCHY | pBL1 | pXYSq10 with full-length linkers | GA | NA |

SOE: Splicing by Overlap Extension;

CPEC: Circular Polymerase Extension Cloning;

GA: Gibson Assembly;

GGA: Golden Gate Assembly;

p1: lacIq promoter.

*: The EcoFlex kit was a gift from Paul Freemont (Addgene kit #1000000080).

**: pDusk was a gift from Andreas Möglich (Addgene plasmid # 43795 ; http://n2t.net/addgene:43795 ; RRID:Addgene_43795)

**Supplementary Table 6. Plasmid assembly detailed descriptions.** Level 2 Golden Gate assembled plasmids are listed in Supplementary Table 4.

| **Plasmid** | **Assembly Method** | **Part** | | |
| --- | --- | --- | --- | --- |
|  |  | **Template / Oligo** | **Forward primer** | **Reverse primer** |
| pXYSj11 | SOE | pBP-lacZ | jXYS31 | jXYS32 |
|  |  | pXYSq2 | jXYS4 | jXYS33 |
| pXYSj15 | SOE | pBP-lacZ | jXYS39 | jXYS43 |
|  |  | gXYS011 | jXYS42 | jXYS44 |
| pXYSq2 | QuikChange | pDusk-p1 (from Childers Lab) | otXYS26 | otXYS27 |
| pXYSq10 | QuikChange | pDusk-p3 (from Childers Lab) | otXYS48 | otXYS49 |
| pXYSq26-C2 | QuikChange | pXYSq2 | otXYS68  otXYS70  otXYS72 | otXYS69  otXYS71  otXYS73 |
| pXYSq28 | QuikChange | pXYSq26-C2 | otXYS66  otXYS68 | otXYS67  otXYS69 |
| pXYSq27-B | QuikChange | pXYSq28 | otXYS64  otXYS74 | otXYS65  otXYS75 |
| pXYSj23 | Gibson | pBP-ORF | jXYS64 | jXYS65 |
|  |  | pXYSq27-B | jXYS66 | jXYS67 |
| pXYSB2 | NEBridge | pT55 | XYS265 | XYS266 |
|  |  | XYS267 | NA | NA |
| pXYSj32 | Gibson | pXYSq27-B | jXYS80 | jXYS81 |
|  |  | pTU2-a-RFP | jXYS82 | jXYS86 |
|  |  | pXYSq27-B | jXYS84 | jXYS85 |
| pXYSj31 | Gibson | pXYSq27-B | jXYS80 | jXYS81 |
|  |  | pTU2-b-RFP | jXYS82 | jXYS83 |
|  |  | pXYSq27-B | jXYS84 | jXYS85 |
| pXYSj43 | Gibson | XYS | jXYS80 | jXYS101 |
|  |  | pTU2-b-RFP | jXYS82 | jXYS102 |
|  |  | pTU2-b-RFP | jXYS103 | jXYS104 |
| pXYSqj43 | QuikChange | pXYSj43 | otXYS80 | otXYS81 |
| pBL1 | Gibson | pXYSq10 | BL1 | BL2 |

**Supplementary Table 7. Gene blocks used in this study.**

| **Name** | **Description** | **Sequence** | **Source** |
| --- | --- | --- | --- |
| gXYS011 | flanking-Start-LZ-GS | ggccacggggcctaccaccatacccacgccgaaacaagcgctcatgagcccgaagtggAgagcccgatcttccccatcgaaaaagagtattgacttcgcatctttttgtacctataatgtgtggagggcccaagttcacttaaaaaggagatcaacaatgaaagcaattttcgtactgaaacatcttaatcatgcacaggagactttctaatgaagcaactcgaagacaaggtcgaagaactcctctcgaagaactaccacctcgagaatgaggtcgcccgtctgggagggggtggctcc | Twist |

**Supplementary Table 8. DNA oligos used in this study.**

| **Name** | **Sequence (5’ → 3’)** |
| --- | --- |
| jXYS4 | CGCATAGGTCTCACTATGACACCATCGAATGGTGCAAAACCTTTCGC |
| jXYS31 | ATCATAAGAGACCCATGCCGTCTCATTAGCTGCAGTCCG |
| jXYS32 | CCATTCGATGGTGTCATAGTGAGACCTATGCGTCTCTAGATTCTAGAAGCGGCCG |
| jXYS33 | GCTAATGAGACGGCATGGGTCTCTTATGATTCACCACCCTCAATTGACTCTCTTCCGG |
| jXYS39 | CCCATAAGAGACCCATGCCGTCTCATTAGCTGCAGTCCG |
| jXYS42 | GCTAATGAGACGGCATGGGTCTCTTATGGGAGCCACCCCCTCCCAGAC |
| jXYS43 | CCTTGTCTTCGAGTTGCTTCATTTTATGAGACCTATGCGTCTCTAGATTCTAGAAGCGGCCG |
| jXYS44 | CGCATAGGTCTCATAAAATGAAGCAACTCGAAGACAAGGTCG |
| jXYS64 | TGCTCAACGATTGAGGATCCTCGAAGAGACCCATGCC |
| jXYS65 | TGTCCCTTGGTCGTCATATGTGAGACCTATGCGTCTCTAGATTCT |
| jXYS66 | GGTCTCACATATGACGACCAAGGGACATATCTACGTCATCG |
| jXYS67 | GGTCTCTTCGAGGATCCTCAATCGTTGAGCATGCCGGCG |
| jXYS80 | GGATCACGGGCGTCCCACGGGTGCGCATGATCG |
| jXYS81 | TCAAAGGATCTTCAACACCCCTTGTATTACTGTTTATGTAAGCAG |
| jXYS82 | CAGTAATACAAGGGGTGTTGAAGATCCTTTGATCTTTTCTACGGGGTC |
| jXYS83 | GCACTTTTCGGGGAAATGTGCTGCAGGGTCTCTGTACCTTCTGAGACG |
| jXYS84 | AGAGACCCTGCAGCACATTTCCCCGAAAAGTGCCAC |
| jXYS85 | GCGCACCCGTGGGACGCCCGTGATCCTGATC |
| jXYS86 | GCACTTTTCGGGGAAATGTGCTGCAGGGTCTCTGTACCCGG |
| jXYS101 | GATCAAAGGATCTTCAACACCCCTTGTATTACTGTTTATGTAAGCAG |
| jXYS102 | GCGTATTGCGTCTCTCCGGATAGTGAGACCTCTAGAAGCGGCCGC |
| jXYS103 | GTCTCACTATCCGGAGAGACGCAATACGCAAACCGCCT |
| jXYS104 | CGGGGAAATGTGCTGCAGGGTCTCTGTACCTTCTGAGACG |
| otXYS26 | CGCCCTTGCTCACCATTAAGCTTGTCGACGG |
| otXYS27 | CCGTCGACAAGCTTAATGGTGAGCAAGGGCG |
| otXYS48 | CTTGCTCACCATAAAGCTTGTCGACGGAG |
| otXYS49 | CTCCGTCGACAAGCTTTATGGTGAGCAAG |
| otXYS64 | GCTTTGTTCAAATGACCGGCTACGAAACCGAGGAAATTT |
| otXYS65 | AAATTTCCTCGGTTTCGTAGCCGGTCATTTGAACAAAGC |
| otXYS66 | CCGAGCTCGTCCATGTCTCCAGGCTGA |
| otXYS67 | TCAGCCTGGAGACATGGACGAGCTCGG |
| otXYS68 | GCCCTGCCGGGTCTATCCTTCGGCT |
| otXYS69 | AGCCGAAGGATAGACCCGGCAGGGC |
| otXYS70 | CCTTCGGCTGCGTAGTCTCCGACGTGC |
| otXYS71 | GCACGTCGGAGACTACGCAGCCGAAGG |
| otXYS72 | CATGCAGCTCCCGAAGACGGTCACAGC |
| otXYS73 | GCTGTGACCGTCTTCGGGAGCTGCATG |
| otXYS74 | GCGATCGCGTATTTCGTCTAGCTCAGGCGC |
| otXYS75 | GCGCCTGAGCTAGACGAAATACGCGATCGC |
| otXYS80 | CGGCCGCTTCTAGAGGTCTCACTATTGCCAGAGACGCAATAC |
| otXYS81 | GTATTGCGTCTCTGGCAATAGTGAGACCTCTAGAAGCGGCCG |
| otXYS82 | GCCTGCTGCCGATGAGATGATAGGAGGTCTAGCATGACGACCAAGG |
| otXYS83 | CCTTGGTCGTCATGCTAGACCTCCTATCATCTCATCGGCAGCAGGC |
| otXYS84 | CGCCAGGGTTTTCCCAG |
| otXYS85 | AGGAAACAGCTATGACCATGATTACG |
| XYS265 | CAAGAATTACAAAGTGAACTTGTTCATGTTTCG |
| XYS266 | TTCGGTAATATCGCGGCTAATCG |
| XYS267 | GGCGATTAGCCGCGATATTACCGAACGCAAAGAACGCGAACAAGAATTACAAAGTGAACTTGTTC |
| BL1 | GCAGCAGAATGAACAGTTGGAGCAGTTCGCAAGTGTCGTTTCTGATCTGACCGAGCACCAGCAGACCCAG |
| BL2 | CCAACTGTTCATTCTGCTGCTCTAACTCCTGCTCACGTTCCTTACGTTCGGTAATATCGCGGCTAATCGCC |

All were ordered from IDT.

**Supplementary Table 9. *E. coli* strains used in this study.** Information on plasmids is listed in Supplementary Table 3-5.

| **Strain ID** | **Plasmid** | **Host Strain** |
| --- | --- | --- |
| WSC2031 | pXYSG2-33 | Top10 |
| WSC1996 | pXYSG2-37 |  |
| WSC2005 | pXYSG2-72 |  |
| WSC1995 | pXYSG2-68 |  |
| WSC2022 | pXYSG2-132 |  |
| WSC2035 | pXYSG2-136 |  |
| WSC2027 | pXYSG2-137 |  |
| WSC2074 | pXYSG2-195 |  |
| WSC2075 | pXYSG2-189 |  |
| WSC2076 | pXYSG2-196 |  |
| WSC2077 | pXYSG2-193 |  |
| WSC2024 | pXYSG2-114 |  |
| WSC2078 | pXYSG2-190 |  |
| WSC2079 | pXYSG2-183 |  |
| WSC2080 | pXYSG2-181 |  |
| WSC2081 | pXYSG2-182 |  |
| WSC2040 | None |  |
| WSC2082 | pXYSG2-198 |  |
| WSC2019 | pXYSG2-111 |  |
| WSC2016 | pXYSG2-108 |  |
| WSC1987 | pXYSG2-70 |  |
| WSC2023 | pXYSG2-112 |  |
| WSC2042 | pXYSG2-174 |  |
| WSC2032 | pXYSG2-76 |  |
| WSC2000 | pXYSG2-150 |  |
| WSC2043 | pXYSG2-128 |  |
| WSC1999 | pXYSG2-149 |  |
| WSC2001 | pXYSG2-151 |  |
| WSC2039 | pXYSG2-159 |  |
| WSC2002 | pXYSG2-147 |  |
| WSC2083 | pXYSG2-197 |  |
| WSC2004 | pXYSG2-67 |  |
| WSC2003 | pXYSG2-89 |  |
| WSC2046 | pXYSG2-173 |  |
| WSC2047 | pXYSG2-172 |  |
| WSC2084 | pXYSG2-194 |  |
| WSC2085 | pXYSG2-187 |  |
| WSC2086 | pXYSG2-192 |  |
| WSC2053 | pXYSjXYS11 |  |
| WSC1941 | pXYSjXYS15 |  |
| WSC1942 | pXYST78 |  |
| WSC1943 | pXYSj23 |  |
| WSC1944 | pXYST23 |  |
| WSC1945 | pXYST55 |  |
| WSC1946 | pXYST24 |  |
| WSC2054 | pXYST79 |  |
| WSC1947 | pXYST28 |  |
| WSC1948 | pXYST47 |  |
| EcoFlex B3 | pBP-lacZ | JM109 |
| EcoFlex B4 | pBP-ORF |  |
| EcoFlex D6 | pBP-SJM914 |  |
| EcoFlex C6 | pBP-J23100 |  |
| EcoFlex C9 | pBP-SJM901 |  |
| EcoFlex D5 | pBP-SJM912 |  |
| EcoFlex C10 | pBP-SJM902 |  |
| EcoFlex D7 | pBP-SJM915 |  |
| EcoFlex D3 | pBP-SJM910 |  |
| EcoFlex D12 | pBP-PET |  |
| EcoFlex E11 | pBP-TL8 |  |
| EcoFlex F1 | pBP-TL10 |  |
| EcoFlex F6 | pBP-L3S2P21 |  |
| EcoFlex G2 | pBP-L2U2H09 |  |
| EcoFlex F5 | pBP-BBa_B0015 |  |
| EcoFlex A2 | pTU1-A-lacZ |  |
| EcoFlex A3 | pTU1-B-lacZ |  |
| EcoFlex A4 | pTU1-C-lacZ |  |
| WSC1955 | pXYSj32 | Top10 |
| WSC1956 | pXYSj31 |  |
| WSC2055 | pXYSqj43 |  |
| WSC1957 | pXYSG1-80 |  |
| WSC2056 | pXYSG1-98-b |  |
| WSC1960 | pXYSG1-131 |  |
| WSC1958 | pXYSG1-129 |  |
| WSC2087 | pXYSG1-151 |  |
| WSC2088 | pXYSG1-192 |  |
| WSC2065 | pXYSG1-146 |  |
| WSC2089 | pXYSG1-167 |  |
| WSC2090 | pXYSG1-148 |  |
| WSC2091 | pXYSG1-171 |  |
| WSC2061 | pXYSG1-97 |  |
| WSC2092 | pXYSG1-165 |  |
| WSC1981 | pXYSG1-11 |  |
| WSC1985 | pXYSG1-138 |  |
| WSC1982 | pXYSG1-135 |  |
| WSC1984 | pXYSG1-137 |  |
| WSC2064 | pXYSG1-139 |  |
| WSC2069 | pXYSG1-156 |  |
| WSC2062 | pXYSG1-120 |  |
| WSC1994 | pXYSq2 |  |
| WSC2059 | pXYSq26-C2 |  |
| WSC2060 | pXYSq27-B |  |
| WSC2093 | pBL1 | Stellar |

**Supplementary Table 10. Fold changes for the circuits.**

| **Figure** | **Plasmid ID** | $\frac{\mathbf{Dark, None}}{\mathbf{Blue light, None}}$ | $\frac{\mathbf{Dark, I3A}}{\mathbf{Blue light,I3A}}$ | $\frac{\mathbf{Dark, None}}{\mathbf{Dark,I3A}}$ | $\frac{\mathbf{Blue light, None}}{\mathbf{Blue light, I3A}}$ | $\frac{\mathbf{Dark, None}}{\mathbf{Blue light, I3A}}$ |
| --- | --- | --- | --- | --- | --- | --- |
| 1c-1 | pXYSG2-33 | 11 |  |  |  |  |
| 1c-2 | pXYSG2-37 |  |  | 4.0 |  |  |
| 1c-3 | pXYSG2-72 | 3.2 | 2.0 | 3.2 | 2.0 | 6.6 |
| 2b-1 | pXYSG2-68 | 12 |  |  |  |  |
| 2b-2 | pXYSG2-132 |  |  | 8.8 |  |  |
| 2b-3 | pXYSG2-136 | 1.9 | 4.1 | 2.1 | 4.4 | 8.5 |
| 3c-1 | pXYSG2-137 | 4.5 |  |  |  |  |
| 3c-2 | pXYSG2-195 |  |  | 15 |  |  |
| 3c-3 | pXYSG2-189 | 1.1 | 5.3 | 1.2 | 5.9 | 6.5 |
| 3c-4 | pXYSG2-196 | 1.3 | 1.0 | 11 | 8.7 | 12 |
| 3c-5 | pXYSG2-193 | 3.3 | 5.6 | 1.1 | 1.9 | 6.4 |
| 4b-1 | pXYSG2-114 | *7.2* |  |  |  |  |
| 4b-2 | pXYSG2-190 |  |  | *6.6* |  |  |
| 4b-3 | pXYSG2-183 | *2.4* | *1.5* | *2.2* | *1.3* | *3.2* |
| 4b-4 | pXYSG2-181 | *1.3* | *2.0* | 4.86 | 3.33 | 2.50 |
| 4b-5 | pXYSG2-182 | 4.0 | 2.2 | *1.1* | *2.1* | 1.94 |
| S1a | None | 0.43 | 0.64 | 0.42 | 0.62 | 0.27 |
| S1b | pXYSG2-198 | 9.9 | 9.1 | 1.4 | 1.3 | 13 |
| S2a-1 | pXYSG2-111 |  |  | 2.2 |  |  |
| S2a-2 | pXYSG2-108 |  |  | 12 |  |  |
| S2a-3 | pXYSG2-110 |  |  | 4.2 |  |  |
| S2a-4 | pXYSG2-37  (same data as in Fig. 1c) |  |  | 4.0 |  |  |
| S2a-5 | pXYSG2-174 |  |  | 1.2 |  |  |
| S2a-6 | pXYSG2-76 |  |  | 1.0 |  |  |
| S2c-1 | pXYSG2-150 | 3.3 | 1.4 | 7.7 | 3.2 | 10 |
| S2c-2 | pXYSG2-128 | 2.7 | 3.3 | 5.6 | 6.9 | 18 |
| S2c-3 | pXYSG2-149 | 1.7 | 2.1 | 3.9 | 4.9 | 8.3 |
| S2c-4 | pXYSG2-70 | 3.3 | 2.6 | 3.2 | 2.5 | 8.4 |
| S2c-6 | pXYSG2-159 | 1.0 | 1.0 | 1.1 | 1.0 | 1.1 |
| S2c-7 | pXYSG2-147 | 0.89 | 0.83 | 1.1 | 1.0 | 0.91 |
| S3b-1 | pXYSG2-197 | 1.7 |  |  |  |  |
| S3b-2 | pXYSG2-67 |  |  | 9.5 |  |  |
| S3b-3 | pXYSG2-89 | 1.7 | 1.3 | 4.7 | 3.6 | 6.1 |
| S3b-4 | pXYSG2-70  (same data as in Fig. S2c) | 3.3 | 2.6 | 3.2 | 2.5 | 8.4 |
| S3b-5 | pXYSG2-173 | 1.3 | 1.4 | 8.8 | 9.3 | 12 |
| S3b-6 | pXYSG2-172 | 1.1 | 0.92 | 14 | 12 | 13 |
| S5-1 | pXYSG2-208 | 1.03 |  |  |  |  |
| S5-2 | pXYSG2-194 |  |  | 1.2 |  |  |
| S5-3 | pXYSG2-187 | 1.0 | 1.8 | 1.0 | 1.9 | 1.9 |

**Supplementary Table 11. Default mathematical modeling parameters for the 2HK-1RR system.**

| **Parameter** | **Value used** | **Description** |
| --- | --- | --- |
| [HK]_T0_ | 0.017 µM | Initial total concentration* of HK |
| [RR]_T0_ | 6 µM | Initial total concentration* of RR |
| *k*_p_ | 0.05 s^-1^ | Dephosphorylation rate constant |
| *k*_a_^+^ in the K state | 0.01-0.3 s^-1^ | HK autophosphorylation rate constant in the K (kinase) state |
| *k*_a_^+^ in the P state | 0.05 s^-1^ | HK autophosphorylation rate constant in the P (phosphatase) state |
| *k*_a_^-^ | 0.001 s^-1^ | HK auto-dephosphorylation rate constant |
| *k*_t_ | 1.5 s^-1^ | Phospho-transfer rate constant |
| *k*_1_^+^ | 1 µM ^-1^ · s^-1^ | Rate constant of the binding of HKP and RR |
| *k*_1_^-^ | 0.5 s^-1^ | Rate constant of the dissociation of HKP and RR |
| *k*_2_^+^ | 1 µM ^-1^ · s^-1^ | Rate constant of the binding of HK and RRP |
| *k*_2_^-^ | 0.5 s^-1^ | Rate constant of the dissociation of HK and RRP |
| *k*_3_^+^ | 1 µM ^-1^ · s^-1^ | Rate constant of the binding of HK and RR |
| *k*_3_^-^ | 0.5 s^-1^ | Rate constant of the dissociation of HK and RR |

The parameters were adapted from Butcher et al.^2^.

At Time 0, [HK] = [HKP] = 0.085 µM; [RR] = [RRP] = 3 µM.

*Definition of “total concentration”:

[HK]_T_=[HK]+[HKP]+[(HK)(RRP)]+[(HKP)(RR)]+[(HK)(RR)]

[RR]_T_=[RR]+[RRP]+[(HK)(RRP)]+[(HKP)(RR)]+[(HK)(RR)]
